# Structure of bacterial phospholipid transporter MlaFEDB with substrate bound

**DOI:** 10.1101/2020.06.02.129247

**Authors:** Nicolas Coudray, Georgia L. Isom, Mark R. MacRae, Mariyah N. Saiduddin, Gira Bhabha, Damian C. Ekiert

## Abstract

In double-membraned bacteria, phospholipids must be transported across the cell envelope to maintain the outer membrane barrier, which plays a key role in antibiotic resistance and pathogen virulence. The Mla system has been implicated in phospholipid trafficking and outer membrane integrity, and includes an ABC transporter complex, MlaFEDB. The transmembrane subunit, MlaE, has minimal sequence similarity to other ABC transporters, and the structure of the entire inner membrane MlaFEDB complex remains unknown. Here we report the cryo-EM structure of the MlaFEDB complex at 3.05 Å resolution. Our structure reveals that while MlaE has many distinct features, it is distantly related to the LPS and MacAB transporters, as well as the eukaryotic ABCA/ABCG families. MlaE adopts an outward-open conformation, resulting in a continuous pathway for phospholipid transport from the MlaE substrate-binding site to the pore formed by the ring of MlaD. Unexpectedly, two phospholipids are bound in the substrate-binding pocket of MlaFEDB, raising the possibility that multiple lipid substrates may be translocated each transport cycle. Site-specific crosslinking confirms that lipids bind in this pocket *in vivo*. Our structure provides mechanistic insight into substrate recognition and transport by the MlaFEDB complex.

## Introduction

The bacterial outer membrane (OM) is a critical barrier that protects the cell from antibiotics and other environmental threats, and protects pathogenic bacteria from the anti-microbial responses of the host. The OM is an asymmetric bilayer, with an outer leaflet of lipopolysaccharide (LPS) and a phospholipid inner leaflet. The OM is separated from the inner membrane (IM) by the periplasmic space, which contains the peptidoglycan cell wall. While this complex envelope architecture has many advantages, it also presents many challenges for OM assembly and transport, including the need to move cargo across two lipid bilayers. Moreover, energy from ATP and the proton motive force are associated with the cytoplasm and inner membrane (IM), leaving the periplasm and OM without direct access to these conventional energy sources. Consequently, double membraned bacteria have evolved a fascinating array of protein machines to overcome the challenge of transporting molecules beyond the IM. These include passive catalysts for OM protein insertion (BAM complex (Knowles *et al.*, 2009; Gu *et al.*, 2016; Han *et al.*, 2016)), as well as ATP and proton-driven machines that couple energy from the cell interior to transport across the cell envelope. An elegant example of this coupling is illustrated by the LPS transport system, which couples an IM ABC (<u>A</u>TP <u>b</u>inding <u>c</u>assette) transporter to a periplasmic bridge and OM complex to export newly synthesized LPS from the IM to the outer leaflet of the OM (Okuda, Freinkman and Kahne, 2012; Okuda *et al.*, 2016; Sperandeo, Martorana and Polissi, 2017). In contrast to the trafficking and assembly of proteins and LPS in the OM, we know comparatively little about how phospholipids are trafficked and inserted into the inner leaflet of the OM, or how the asymmetry of the OM is maintained.

The Mla system, an ABC transporter in *E. coli* and related Gram-negative bacteria, has recently emerged as a key player in phospholipid transport across the bacterial envelope. Mla trafficks phospholipids between the IM and OM and is important for maintaining the outer membrane barrier (Malinverni and Silhavy, 2009; Chong, Woo and Chng, 2015; Thong *et al.*, 2016; Abellón-Ruiz *et al.*, 2017; Ekiert *et al.*, 2017; Isom *et al.*, 2017; Shrivastava, Jiang and Chng, 2017; Powers and Trent, 2018; Yeow *et al.*, 2018; Ercan *et al.*, 2019; Hughes *et al.*, 2019; Kamischke *et al.*, 2019; Shrivastava and Chng, 2019). This system consists of three main parts: 1) an IM ABC transporter complex, MlaFEDB; 2) an OM complex, MlaA-OmpC/F; and 3) a soluble periplasmic protein, MlaC, which has been proposed to shuttle phospholipids between MlaFEDB and MlaA-OmpC/F (**Figure 1a**). The directionality of transport facilitated by the Mla pathway is still an area of intense research, with reports of both phospholipid import (Malinverni and Silhavy, 2009; Chong, Woo and Chng, 2015; Powers and Trent, 2018; Yeow *et al.*, 2018) and export (Hughes *et al.*, 2019; Kamischke *et al.*, 2019). The IM complex, MlaFEDB, consists of four different proteins: MlaD, a membrane anchored protein from the MCE (<u>M</u>ammalian <u>C</u>ell <u>E</u>ntry) protein family, which forms a homohexameric ring in the periplasm (Thong *et al.*, 2016; Ekiert *et al.*, 2017); MlaE (also called YrbE), a predicted integral inner membrane ABC permease; MlaF, an ABC ATPase; and MlaB, a STAS (<u>S</u>ulfate <u>T</u>ransporter and <u>A</u>nti-<u>S</u>igma factor antagonist) domain protein with possible regulatory function (Kolich *et al.*, 2020). Crystal structures of MlaD (Ekiert *et al.*, 2017) and MlaFB (Kolich *et al.*, 2020) have provided insights into the function of individual domains and transporter regulation, but the structure of the transmembrane subunit, MlaE, has been lacking. MlaE has no detectable sequence similarity to proteins of known structure or function, suggesting it adopts a unique or divergent ABC transporter fold. Low resolution cryo electron microscopy (cryo-EM) studies (Ekiert *et al.*, 2017; Kamischke *et al.*, 2019) have established the overall shape of the complex, but have not shed much light on how the various subunits of the MlaFEDB complex assemble and function. Thus, a structure of the MlaFEDB complex may provide important insights into the mechanisms of bacterial lipid transport, as well as the evolution and function of the MlaE/YrbE transporters, which are conserved from double-membraned bacteria to chloroplasts.

**Figure 1.**
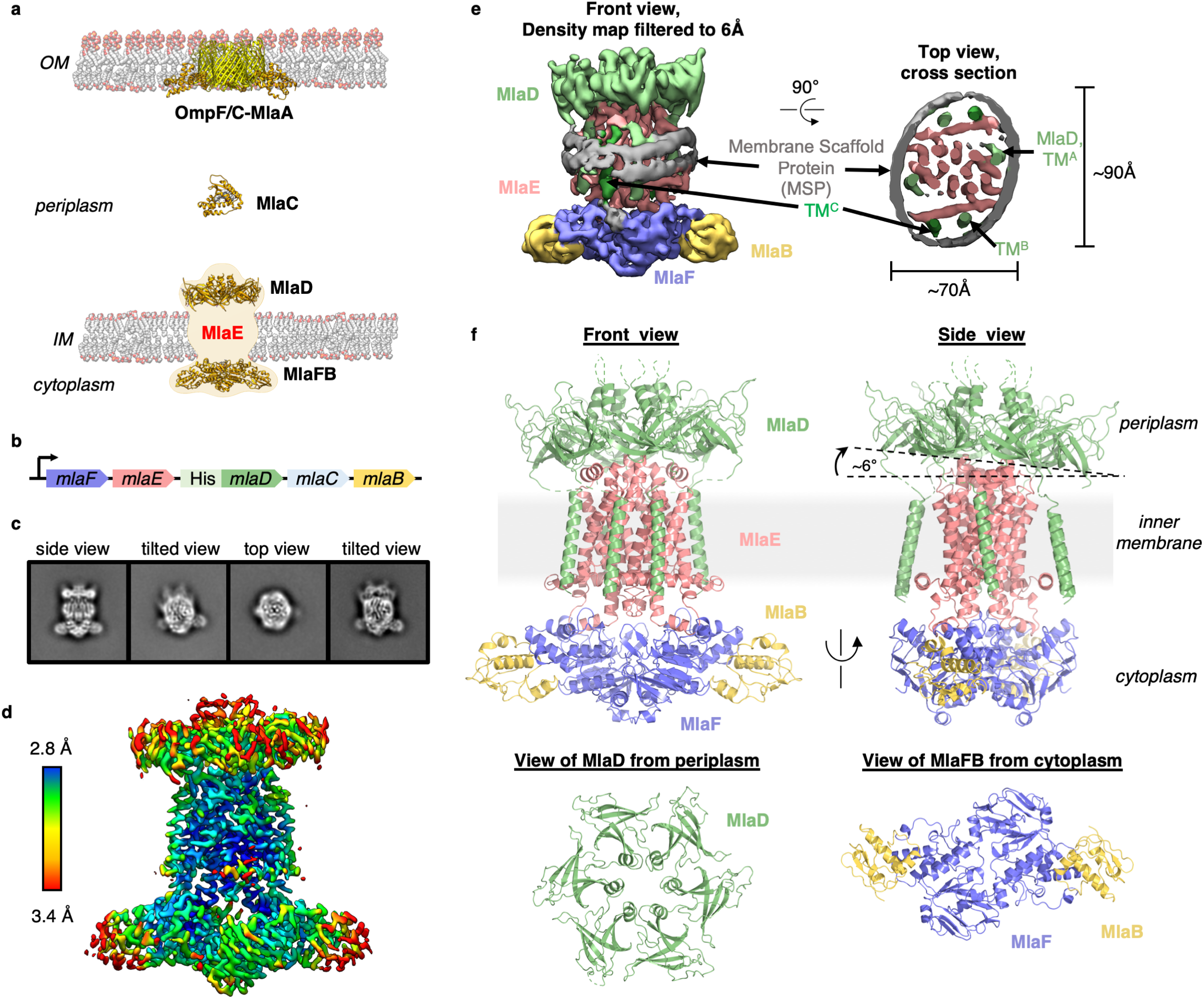
Cryo-EM structure of the MlaFEDB complex. **(a)** Schematic of the Mla pathway (adapted from (Kolich *et al.*, 2020)). The OmpF/C-MlaA complex (PDB 5NUP), periplasmic shuttle protein MlaC (PDB 5UWA), and MlaFEDB complex (PDB 5UW2 and 6XGY, EMDB-8610) are shown. **(b)** Schematic of the MlaFEDCB operon with N-terminal His-tag on MlaD, as reported previously (Ekiert *et al.*, 2017). **(c)** 2D class averages from single particle cryo-EM analysis of MlaFEDB in nanodiscs. **(d)** Final EM density map of MlaFEDB, colored by local resolution (EMD-22116). **(e)** Density map of MlaFEDB filtered to 6 Å, showing membrane scaffold protein (MSP) belts surrounding the edge of the lipid nanodisc. MlaF, slate blue; MlaE, pink; MlaD, green; MlaB, yellow; MSP, grey. The packing of the 6 TM helices from the MlaD subunits around the periphery of the MlaE core is apparent in the Top View. **(f)** Overview of the MlaF_2_E_2_D_6_B_2_ model (PDB 6XBD); colors as in Figure 1e. Regions of disorder in MlaD linkers and C-termini are indicated by green dashed lines. The MlaD ring is tilted relative to MlaFEB, resulting in a complex that is asymmetric overall.

## Results

### Overview of the MlaFEDB structure

To address how Mla drives lipid transport, we overexpressed the *mla* operon (**Figure 1b**) and reconstituted the MlaFEDB ABC transporter complex in lipid nanodiscs containing *E. coli* polar lipids (see Methods), and determined the structure using single particle cryo-EM, with an average resolution of 3.05 Å (**Figure 1c-e**, **Figure 1 — figure supplement 1 and 2, Supplementary file 1)**. Although MlaFEDB was expected to exhibit 2-fold symmetry, initial reconstructions showed clear asymmetry in our maps, which we then refined without applying symmetry (**Figure 1d, Figure 1 — figure supplement 1 and 2, Supplementary file 2**). Local resolution analysis showed that the entire complex is well-defined at ~2.8 – 3.5 Å resolution (**Figure 1d, Figure 1 — figure supplement 3a**), allowing us to build a nearly complete model for MlaFEDB (**Figure 1f**), including a high-resolution structure of the MlaE transmembrane subunit. In addition, we resolved both coils of the membrane scaffold protein (MSP) belt surrounding the nanodisc (Bayburt, Grinkova and Sligar, 2002) using the map filtered at 6Å, thereby clearly defining the position of the transmembrane domain (**Figure 1e, Figure 1 — figure supplement 3a,b**).

The MlaFEDB transporter is significantly larger and more complicated than most ABC transporter structures determined to date, consisting of a total of 12 polypeptide chains from 4 different genes, with a stoichiometry of MlaF_2_E_2_D_6_B_2_. At the center of the complex, a core ABC transporter module is formed from two copies each of the MlaF ATPase and the MlaE transmembrane domains (TMDs). Outside this ABC transporter core, MlaFEDB contains additional subunits not found in other ABC transporters: MlaD on the periplasmic side of the IM, and MlaB in the cytoplasm. A homohexameric ring of MCE domains from MlaD sits atop the periplasmic end of MlaE like a crown. This MCE ring is anchored in place by six MlaD transmembrane helices, which dock around the periphery of the MlaE TMDs (**Figure 1f**). On the cytoplasmic side, each of the MlaF ATPase subunits is bound to MlaB, a STAS domain protein that was recently reported to act as regulator of the MlaFEDB transporter (Kolich *et al.*, 2020). The overall structure of the MlaF_2_B_2_ module is very similar to our recent MlaFB X-ray structure (PDB: 6XGY), apart from a small relative rotation between the MlaF helical and catalytic subdomains (**Figure 1 — figure supplement 4a**). This rotation is similar to motions described in other ABC transporters (Karpowich *et al.*, 2001; Smith *et al.*, 2002; Orelle *et al.*, 2010). An unusual C-terminal extension of each MlaF protomer wraps around the neighboring MlaF subunit and docks near the MlaFB interface, almost identical to the domain-swapped “handshake” motif observed in the crystal structure of the MlaF_2_B_2_ subcomplex (**Figure 1 — figure supplement 4b**) (Kolich *et al.*, 2020). While the MlaFEB subcomplex exhibits near-perfect 2-fold rotational symmetry at this resolution, the MlaD ring is clearly tilted relative to MlaE, resulting in a misalignment of the 2-fold symmetry axis of MlaFEB and the pseudo-6-fold axis of MlaD by approximately 6 degrees (**Figure 1f**).

### MlaE is distantly related to the TMDs of other ABC transporters

The transmembrane subunits of ABC transporters play a central role in determining the transport mechanism and substrate specificity. Consequently, the structure of the MlaE subunit is of particular interest. Our cryo-EM structure reveals that the core TMD of MlaE consists of 5 transmembrane helices (TM1 – TM5) (**Figure 2a, 2e-g**). A coupling helix (CH) in the cytoplasm connects TM2 and TM3, and mediates the interaction between the TMDs of MlaE and the MlaF ATPase subunits (**Figure 1 — figure supplement 4c**). A small periplasmic helix (PH) is found between TM3 and TM4 at the periplasmic side of MlaE. Two additional N-terminal helices are membrane embedded, which we call interfacial helices 1 and 2 (IF1 and IF2 (Chen *et al.*, 2020); discussed in more detail below). The IF1 helix is a 30 residue long, amphipathic N-terminal helix that lies parallel to the membrane within the cytoplasmic leaflet, and extends along the width of the MlaE dimer (**Figure 2a, 3b**). IF2 is angled relative to the plane of the membrane, and is separated from TM1 by a kink within the lipid bilayer. While the C-terminal portion of IF1 interacts with TM3 and TM4, the N-terminal half projects outward into the surrounding membrane, creating a cleft between the core TMD and the IF1 helix (**Figure 3a, b**). Additional EM densities were observed in this cleft (**Figure 1 — figure supplement 5**), which may be phospholipids or other molecules; these ligands were not modeled explicitly as their identities are ambiguous.

**Figure 2.**
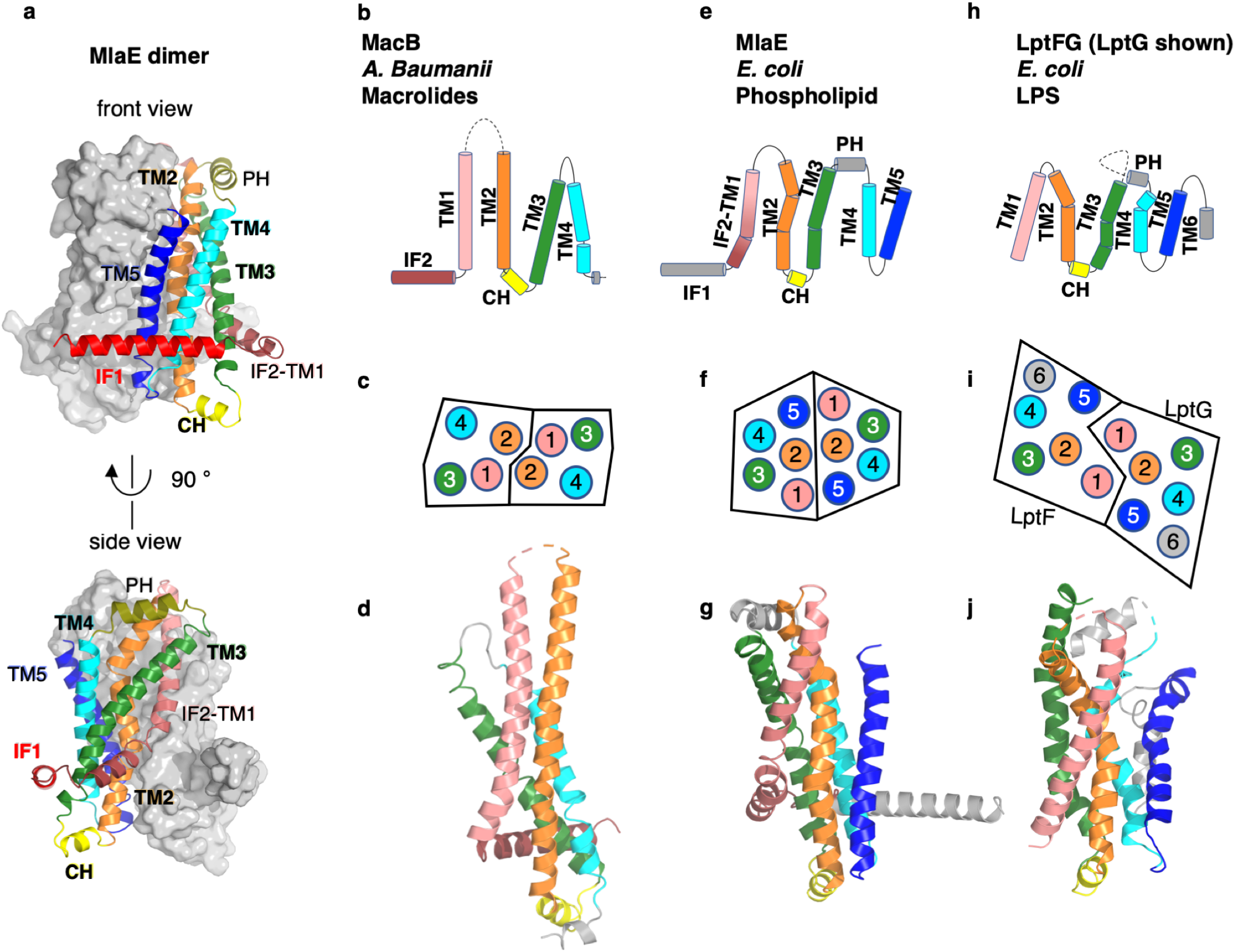
Topology and fold of MlaE. **(a)** MlaE dimer, with one protomer represented as surface, and the other as cartoon. **(b-j)** comparison of MlaE (PDB 6XBD) with the related transmembrane domains of ABC transporters, MacB and LptFG (PDB 5GKO and 6MHZ). **(b, e, h)** Topology diagrams. CH, coupling helix; PH, periplasmic helix; IF, interfacial helix (also called connecting helix in ABCG transporters); TM, transmembrane helix. **(c, f, i)** Schematics representing helices at the dimer interface, viewed from the periplasm (each circle represents a helix). **(d, g, j)** Cartoon view of monomer. See **Figure 2 — figure supplement 1** for comparisons with additional related transporters.

**Figure 3.**
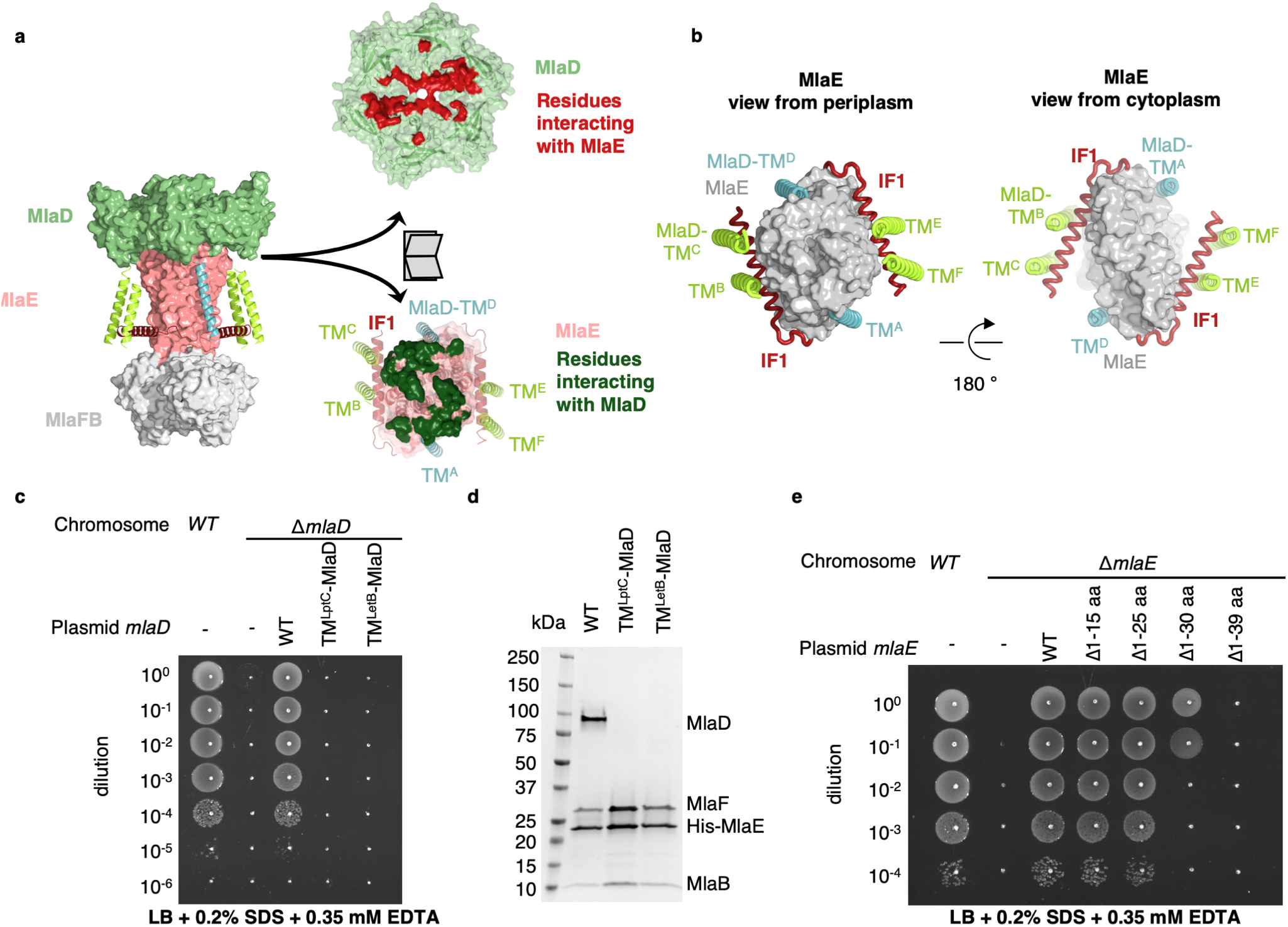
TM helices of MlaD are important for interaction with MlaE. **(a)** Structure of MlaFEDB complex, highlighting interacting regions between MlaE and MlaD. MlaD-TM^A/D^ (cyan helices, cartoon representation) interact with the core domain of MlaE (pink, surface representation); MlaD-TM^B/C/E/F^ (green helices, cartoon representation) interact mostly with IF1 (red helices, cartoon representation). The periplasmic end of the MlaE dimer interacts with the MlaD ring (shown in “open book” representation, right). In “open book” representation, MlaD residues that interact with MlaE are shown in red, and MlaE residues that interact with MlaD are shown in green, as determined using COCOMAPS (Vangone *et al.*, 2011). **(b)** Top and bottom views highlighting the interaction of helices between MlaE and MlaD. Helices colored as in (a). **(c)** Cellular assay for the function of MlaD. 10-fold serial dilutions of the indicated cultures spotted on LB plates containing SDS+EDTA at the concentrations indicated, and incubated overnight. The *mlaD* knockout does not grow in the presence of SDS+EDTA, but can be rescued by the expression of WT MlaD from a plasmid. MlaD chimeras containing LptC or LetB TM helices fail to complement. Corresponding controls plated on LB only can be found in **Figure 3 — figure supplement 2a**. **(d)** SDS-PAGE of recombinantly expressed and purified complexes formed in the presence of WT MlaD or MlaD chimeras containing LptC or LetB TM helices. **(e)** Cellular assay for the function of MlaE, as described for MlaD in (c). Small deletions of the N-terminus of MlaE IF1 are tolerated, while larger deletions impair or completely abolish growth. Corresponding controls plated on LB only can be found in **Figure 3 — figure supplement 2d**.

Despite negligible sequence similarity, the core MlaE fold is related to the TMDs of several other ABC transporters. MlaE most closely resembles the LPS exporter LptF/LptG (Thomas *et al.*, 2020) and the macrolide antibiotic efflux pump MacB (Crow *et al.*, 2017; Fitzpatrick *et al.*, 2017; Murakami, Okada and Yamashita, 2017) (**Fig. 2b-j**), but also shares similarities with the glycolipid flippases Wzm (Bi *et al.*, 2018; Caffalette *et al.*, 2019) and TarG (Chen *et al.*, 2020) and the eukaryotic ABCA/ABCG families (**Figure 2 — figure supplement 1**). However, MlaE also displays notable differences. First, previously determined structures from the above families contained either 4 TM helices (MacB) or 6 TM helices (all the rest). In contrast, MlaE is intermediate between these two groups; MacB contains TM1-TM4, MlaE contains TM1-TM5, and other transporters contain TM1-TM6 (**Figure 2, Figure 2 — figure supplement 1)**. Second, in both MlaE and LptFG, TM5 exists as one continuous helix, whereas TarG, Wzm, and ABCA/ABCG have a small insertion near the periplasmic end that results in a pair of reentrant helices (**Figure 2 — figure supplement 2a**). Third, MlaE has a membrane-embedded helix preceding TM1, which we call IF2 (also called CnH (Lee *et al.*, 2016), IF (Bi *et al.*, 2018), or “elbow helix”). In MacB, TarG, Wzm and ABCA/ABCG transporters, IF2 forms an amphipathic helix that runs roughly parallel to the membrane within the cytoplasmic leaflet, followed by a sharp turn and a separate TM1, which spans nearly the entire bilayer (**Figure 2 — figure supplement 2b**). In contrast, LptFG has no clear IF2 counterpart, as these TMDs begin with TM1 forming a long, continuous helix (**Figure 2 — figure supplement 2b**). In MlaE, this region adopts an intermediate configuration, where IF2 and TM1 are both involved in the first traverse of the membrane, but these two segments are distinguished by a clear kink in the middle of the bilayer (**Figure 2 — figure supplement 2b**). Overall, the structure of the transmembrane region of MlaE differs significantly from previously determined structures, but has at its core a transporter domain conserved across a structurally diverse group of ABC transporters and shared between bacteria and eukaryotes.

### Interactions between MlaE and MlaD

MlaE has three main modes of interaction with MlaD (**Figure 3a**). First, the MlaE core TMDs interact with 2 of the MlaD TM helices; second, the IF1 helices of the MlaE dimer interact with the remaining 4 MlaD TMs; and third, the periplasmic end of the MlaE dimer interacts with the MlaD ring. Due to the symmetry mismatch between the pseudo-6-fold symmetric MlaD hexamer and the 2-fold symmetric MlaFEB module, the six identical transmembrane helices of MlaD interact with MlaE in 3 non-equivalent ways (**Figure 3b**). The MlaD TMs from chains A and D (MlaD-TM^A/D^) are closely packed against the MlaE TMDs on opposite sides of the complex. The remaining 4 TM helices from MlaD are largely isolated in the membrane, and their main interactions are with the MlaE IF1 helices via helix crossing interactions, with MlaD-TM^B/E^ and MlaD-TM^C/F^ contacting IF1 residues 6-14 and 17-25, respectively (~84° crossing angle, **Figure 3a, Figure 3 — figure supplement 1**).

To test whether the MlaD TM helices are important for MlaFEDB complex assembly, we generated chimeras in which we replaced the native TM helix of MlaD with a TM helix expected to make no direct interactions with MlaE [from the *E. coli* IM proteins LptC (TM^LptC^) or LetB (TM^LetB^)]. We tested the ability of these MlaD chimeras to complement an *mlaD* knockout strain of *E. coli*. Mutations in components of the Mla pathway exhibit a substantial growth defect in LB medium in the presence of SDS+EDTA (Malinverni and Silhavy, 2009), which can be complemented with WT *mlaFEDCB* on a plasmid. We found that neither of the MlaD chimeras were able to restore growth of an *mlaD* knockout strain in the presence of SDS+EDTA (**Figure 3c, Figure 3 — figure supplement 2a**). To assess whether these MlaD chimeras are still capable of interacting to form MlaFEDB complexes, we recombinantly overexpressed the *mlaFEDCB* operon encoding either WT or chimeric MlaD, and purified the resulting complexes using an affinity tag on MlaE. SDS-PAGE showed that MlaE co-purified with MlaB, MlaF and WT MlaD, but MlaD chimeras did not co-purify (**Figure 3d**), despite robust expression of hexameric MlaD in the membrane fraction (**Figure 3 — figure supplement 2b**). Thus, the mere presence of MlaD hexamers anchored to the membrane is not sufficient to complement an *mlaD* knock-out, but rather MlaD appears to require its native TM helix in order to assemble and function in complex with MlaFEB. These results suggest that the MlaD TM helix interactions with MlaE drive specificity in the formation of the complex.

While MlaD-TM^A/D^ helices interact intimately with the core MlaE TMD, MlaD-TM^B/E^ and MlaD-TM^C/F^ interact more loosely with MlaE, mostly through IF1. To test whether the MlaD-TM^B/E^ and MlaD-TM^C/F^ interactions are essential for function in cells, we generated a series of truncation mutants of MlaE IF1 with 15 or 25 residues deleted from the N-terminus of IF1 (Δ1-15 aa and Δ1-25 aa), thereby removing the MlaD-TM^C/F^ binding site or both the MlaD-TM^C/F^ and MlaD-TM^B/E^ binding sites, respectively (**Figure 3 — figure supplement 2c**). We used a similar complementation assay to the one described above, but for an *mlaE* knockout, to assess the function of these variants in cells. We observed that both the Δ1-15 aa and Δ1-25 aa mutants fully restored growth of an *mlaE* knockout under these conditions, similar to complementation by the WT operon (**Figure 3e, Figure 3 — figure supplement 2d**). This result is surprising, as the residues of IF1 interacting with the MlaD TM are highly conserved (**Figure 3 — figure supplement 1**). It is possible that future experiments that probe function more specifically may reveal a role for these conserved residues. Interestingly, MlaE mutants with larger deletions only partially restored growth (Δ1-30 aa), or failed to complement (Δ1-39 aa), suggesting that the C-terminus of IF1 and/or the following loop may have an essential function (**Figure 3e**). All the MlaE truncation mutants expressed well, and formed complexes with MlaD, MlaF and MlaB (**Figure 3 — figure supplement 2e**), though we noted that all of the mutants appeared to incorporate less MlaFB into the complex. The significance of MlaFB destabilization is not clear, though binding of MlaFB to MlaE was previously proposed to be weaker than is typically observed for ABC transporters, and reversible association/dissociation of the complex may be a mechanism of MlaFEDB regulation (Kolich *et al.*, 2020). Taken together, our data show that the TM regions of MlaD form specific interactions with MlaE, and that the tight interactions formed by TM^A/D^ are sufficient for complex assembly and function.

### MlaE adopts an outward-open conformation in the apo state

While the vast majority of ABC transporter structures adopt an inward-open conformation in the absence of nucleotide (Gerber *et al.*, 2008; Kadaba *et al.*, 2008; Aller *et al.*, 2009; Manolaridis *et al.*, 2018), our structure of MlaE is in the outward-open state (**Figure 4a**). This uncommon configuration was previously observed in the related transporters LptFG (Li, Orlando and Liao, 2019b; Owens *et al.*, 2019) and MacB (Fitzpatrick *et al.*, 2017), suggesting that the Mla, Lpt and MacAB may share some mechanistic features. The narrow outward-open pocket within MlaE encloses a volume of ~750 Å^3^ (estimated using CASTp (Tian *et al.*, 2018)), and is primarily formed by TM1 and TM2, with some contribution from TM5 (**Figure 4a**). This is similar to LptFG, where TM1, TM2, and TM5 form a much larger pocket (volume of ~3000 Å^3^, estimated using CASTp (Tian *et al.*, 2018), in PDB: 6MHU (Li, Orlando and Liao, 2019a)) at the periplasmic side of the complex for LPS binding. The pockets in both transporters are largely hydrophobic in nature, consistent with binding to lipid substrates, though in LptFG the rim has a pronounced positive charge proposed to interact with phosphates on the LPS inner core (Li, Orlando and Liao, 2019a), while MlaE is more neutral.

**Figure 4.**
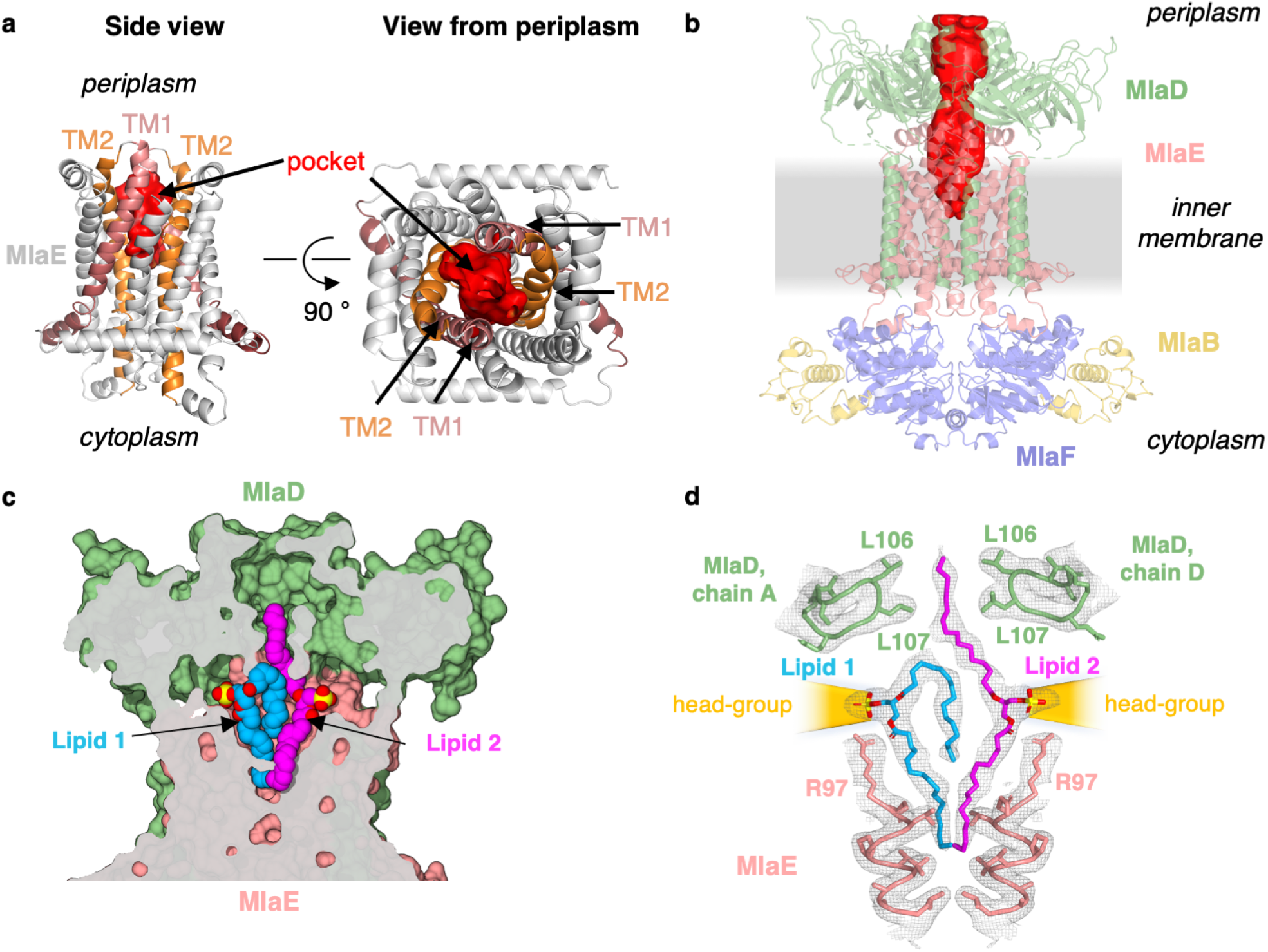
Lipids are bound in an outward-open pocket formed by MlaE and MlaD. **(a)** Side view (left) and view from periplasm (right) of MlaE dimer highlighting the outward-open pocket formed by TM1 (salmon helices) and TM2 (orange helices). The boundary of the substrate-binding pocket was estimated using CASTp (Tian *et al.*, 2018), and is displayed as a red surface. **(b)** Side view of MlaFEDB complex, showing the continuous hydrophobic channel running from the substrate binding pocket in MlaE to the periplasmic space, through the pore of MlaD (red, tunnel boundary estimated with HOLLOW (Ho and Gruswitz, 2008)). **(c)** Side view cross-section of the hydrophobic channel between MlaE and MlaD showing two bound phospholipids in blue and magenta. **(d)** Lipids modeled with surrounding structural elements and key residues highlighted. EM density map is shown as mesh, at the same contour level for lipids and surrounding regions. The lateral channels that could accommodate lipid head groups are indicated by orange cones.

MlaD forms a hexameric MCE ring with a hydrophobic pore at the center (Ekiert *et al.*, 2017). Similar hydrophobic tunnels have been observed through the MCE rings of PqiB and LetB (Ekiert *et al.*, 2017; Isom *et al.*, 2020), and phospholipids have been crosslinked inside the tunnel of LetB. Indeed, in our structure of the MlaFEDB complex, the pore of MlaD and the outward-open pocket of MlaE line up with each other, resulting in a continuous hydrophobic tunnel that runs from the pocket of MlaE, through MlaD, and out into the periplasm (**Figure 4b**). Conceptually similar to the MlaE-MlaD channel, structurally diverse lipid transport domains have also been resolved for several other ABC transporters (Qian *et al.*, 2017; Li, Orlando and Liao, 2019a; Owens *et al.*, 2019) (**Figure 4 — figure supplement 1**), and are proposed to facilitate the movement of hydrophobic molecules through the aqueous extracellular/periplasmic environment.

### Lipids are bound in the substrate-translocation pathway

Additional density was apparent in the outward-open pocket of MlaE, which is the right size and shape to accommodate two diacyl phospholipids **(Figure 4c, d)**. We modeled these as the most abundant PL species in *E. coli*, phosphatidylethanolamine, and these lipids make extensive contacts with both MlaE subunits as well as MlaD **(Figure 4 — figure supplement 2a**; see **Methods** for additional details). Unlike recent structures of the LPS exporter bound to LPS, where all of the acyl chains project downward into the hydrophobic pocket of LptFG (Li, Orlando and Liao, 2019b; Tang *et al.*, 2019), the lipids bound to MlaFEDB appear to be trapped in different conformations (**Figure 4c)**, perhaps intermediates in the process of being transferred between MlaE and the MlaD pore. In one lipid molecule (lipid 1), both acyl chains are bound in the pocket of MlaE (**Figure 4c, d**). Strikingly, the second lipid (lipid 2) adopts an extended conformation, with one acyl chain reaching down into the MlaE pocket while the other projects upwards to insert into a constriction in the MlaD pore formed by Leu106 and Leu107 (**Figure 4d**), which have previously been implicated in MlaD function (Ekiert *et al.*, 2017). The upward-facing acyl chain of lipid 2 is sandwiched between two tyrosine residues (Tyr81 from each MlaE subunit), and these residues also contact one of the acyl chains of lipid 1. Mutations of Tyr81 to either a smaller (Ala) or a larger (Trp) hydrophobic residue had no effect on *E. coli* growth in the presence of SDS+EDTA (**Figure 4 — figure supplement 3)**, though a Tyr81Glu mutation was recently reported to impair MlaFEDB function (Tang *et al.*, 2020) (see **Discussion**).

In contrast to the hydrophobic fatty acid tails, which are completely buried within the MlaE-MlaD tunnel, the polar head group of each lipid projects outwards through lateral solvent accessible channels on opposite sides of the complex. Beyond the phosphoglycerol core, the density for the lipid head groups in the lateral channels of MlaFEDB are not well resolved, and the only noteworthy interaction is a salt bridge formed between Arg97 of MlaE and the head group phosphate (**Figure 4d**). Weaker density beyond the phosphate may reflect heterogeneity in the lipid species bound to MlaFEDB in our structure, and/or that the interactions between the headgroup and nearby MlaE and MlaD residues may be weak and non-specific. The crystal structure of MlaC bound to phospholipid revealed a similar binding mode and lack of head group specificity (Ekiert *et al.*, 2017). Thus, rather than mediate the binding of specific lipids, the lateral channels may instead serve as a non-specific cavity to accommodate a range of polar head groups during the “lipid gymnastics” (Neumann, Rose-Sperling and Hellmich, 2017) that may need to occur to translocate lipids between the MlaE pocket and the MlaD pore, which may involve flipping the lipids upside down (see **Discussion**). In addition to interactions with the lipid head groups, Arg97 is part of a cluster of conserved charged residues, including Glu98, Lys205, and Asp250, which form salt bridges buried in the hydrophobic core of MlaE and are part of a larger polar interaction network including Gln73, Asp198, and Thr254 (**Figure 3 — figure supplement 1c, Figure 4 — figure supplement 3a**). To probe the potential role of these residues in MlaFEDB function, we mutated Arg97, Glu98, Lys205, and Asp250 individually to alanine. Surprisingly, we found these single mutations had no effect on *E. coli* growth in the presence of SDS+EDTA (**Figure 4 — figure supplement 3**; see **Discussion**). Thus, the role of these conserved residues remains unclear.

The presence of two lipid densities in our structure raises the possibility that the Mla system may transport two substrates per transport cycle. Structures of the periplasmic lipid carrier protein, MlaC, have been determined with either 1 or 2 diacyl phospholipids bound (Ekiert *et al.*, 2017) **(Figure 4 — figure supplement 4**), and a structure of apo *E. coli* MlaC revealed a clamshell-like motion resulting in significant changes in the volume of the lipid binding pocket (Hughes *et al.*, 2019). The different architectures and conformational states of the MlaC pocket suggest that it may accommodate one or two phospholipids, or larger lipid molecules. Indeed, previous studies have suggested that cardiolipin may be a substrate of the Mla system (Kamischke *et al.*, 2019), and that cardiolipin is detected by TLC on lipid extracts from components of the Mla system (Hughes *et al.*, 2019). Together with previous functional data, the presence of 2 phospholipids/4 acyl chains bound in our MlaFEDB structure raises the possibility that the Mla system may also be capable of transporting tetra-acyl lipids, such as cardiolipin. We note, however, that cardiolipin bound to MlaE would have to adopt a somewhat different configuration from the lipids observed in our structure, in order to covalently link the two phosphate groups closer together.

Viewed from the side, the MlaD ring is tilted with respect to MlaFEB by approximately 6 degrees (**Figure 1f**), and deviates from the expected 6-fold symmetry observed in the crystal structure of MlaD and other MCE proteins (**Figure 4— figure supplement 2b, c**). Differences between subunits that result in symmetry breaking include re-organization of 2 of the 6 pore lining loops (containing Leu107) as well as domain level rearrangements in the ring (**Figure 4 — figure supplement 2b, c**). This is particularly surprising, as the MlaFEB module is 2-fold symmetric in our EM structure, and the crystal structure of the MlaD ring in isolation exhibited near perfect 6-fold symmetry. This raises the question: what is breaking the symmetry in our MlaFEDB structure? The clear asymmetric density for the two bound phospholipids suggests that the asymmetry of the MlaD ring and the configuration of the lipids is correlated; otherwise, the cryo-EM reconstruction would yield 2-fold symmetric lipid densities to match the 2-fold symmetric features of the MlaFEB module. Thus, the asymmetry in MlaD appears to arise from its asymmetric interactions with lipid 1 at the interface of MlaE and MlaD. Leu107 from MlaD chain F makes hydrophobic interactions with one of the fatty acid tails, perhaps stabilizing this side of the MlaD ring in closer proximity to MlaE and the lipid binding pocket. The resulting conformational changes in the MlaD ring could be important for lipid translocation through the channel or perhaps even modulating the binding of MlaC to the transporter and facilitating lipid transfer between MlaD and MlaC (see Discussion).

To assess whether phospholipids are bound in the pocket of MlaE in cells, we utilized a site-specific photocrosslinking method (Isom *et al.*, 2020) to detect binding of radiolabeled phospholipids *in vivo*. We incorporated the unnatural photocrosslinking amino acid *p*-benzoyl-L-phenylalanine (BPA) (Chin *et al.*, 2002) into the MlaE protein at five positions in the lipid binding site (Leu70, Val77, Leu78, Tyr81, and Leu99), as well as Phe209 (protected in the MlaE core; not expected to contact lipids) or Trp149 (membrane exposed; expected to contact bulk membrane lipids) (**Figure 5a**). After crosslinking in cells grown in the presence of ^32^P orthophosphate to label total phospholipids, these MlaFEDB complexes were purified and analyzed by SDS-PAGE. We observed both a monomeric and dimeric form of MlaE, the latter of which was enriched in crosslinked samples where BPA had been incorporated at the dimer interface (**Figure 5 — figure supplement 1, Figure 5 — figure supplement 2**). However, as the level of dimerization was variable between mutants, we focused only on the monomeric band for our analysis (**Figure 5b)**. Crosslinking at Trp149 and Phe209 resulted in high and low ^32^P signal, respectively, indicative of an abundance of phospholipids near the membrane-exposed Trp149 and very few phospholipids near the buried Phe209, as expected (**Figure 5b**). ^32^P incorporation into MlaE was induced by crosslinkers at all five positions in the outward-open lipid binding pocket, with particularly high signals for Tyr81 and Leu99 (**Figure 5b)**. Furthermore, the uncrosslinked controls showed a weak signal, indicating that the elevated ^32^P signal in the crosslinked samples was due to the formation of crosslinks between BPA and phospholipids at those locations. Thus, the phospholipid binding site observed in our MlaFEDB structure is occupied by phospholipids *in vivo*.

**Figure 5.**
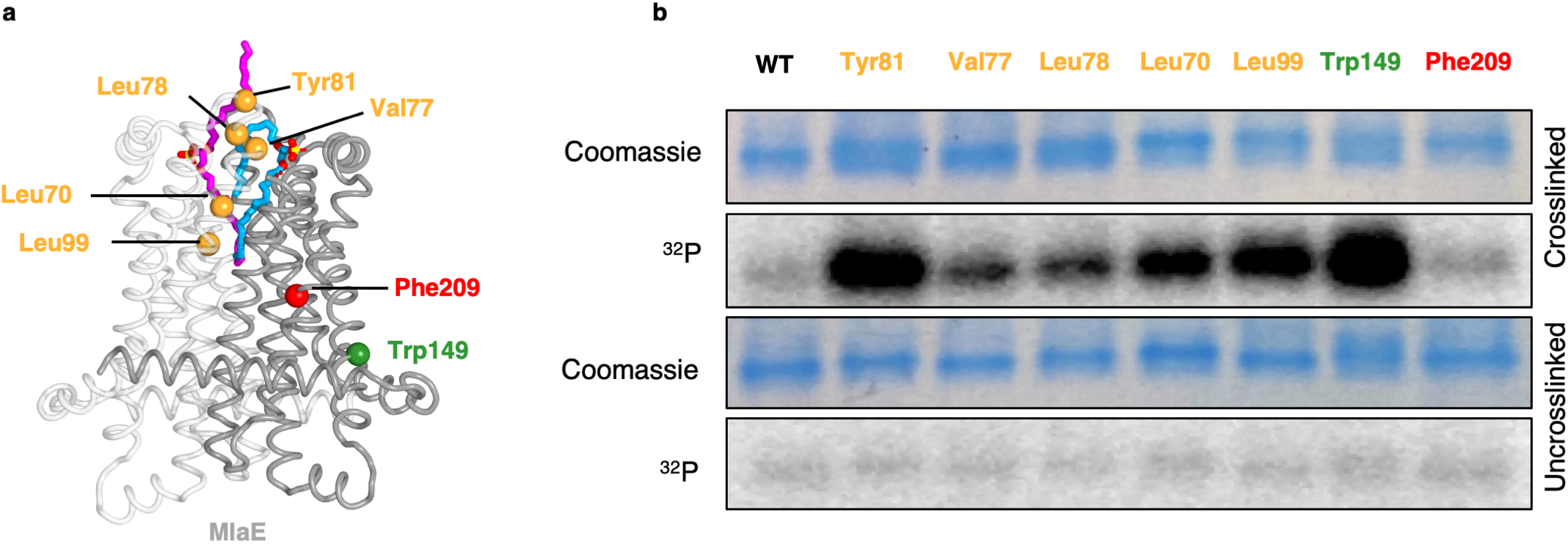
*In vivo* photocrosslinking of substrates in MlaFEDB. **(a)** MlaE dimer (gray cartoon) showing sites of BPA crosslinker incorporation (spheres). Orange, residues in the lipid-binding pocket; red, residue buried within the MlaE core, designed as a negative control; green, residue facing the membrane environment, designed as a positive control. Bound lipids are shown in magenta and blue sticks. **(b)** SDS-PAGE of purified WT MlaFEDB and BPA mutants, either crosslinked or uncrosslinked, and visualized by Coomassie staining (protein) or phosphorimaging (^32^P signal). Band corresponding to the MlaE monomer is shown here, and full gels are shown in **Figure 5 — figure supplement 1**.

## Discussion

The results presented here provide mechanistic insights into lipid transport by the Mla system. Our structure reveals that MlaE is structurally related to two bacterial exporters (LPS exporter and MacAB). A role for MlaFEDB in phospholipid export is supported by recent cellular studies in *Acinetobacter baumannii (Kamischke et al., 2019)*, as well as *in vitro* experiments with *E. coli* proteins, showing directional lipid transfer from MlaD to MlaC (Ercan *et al.*, 2019; Hughes *et al.*, 2019). On the other hand, prior studies indicated that the Mla system is an importer (Malinverni and Silhavy, 2009; Chong, Woo and Chng, 2015; Powers and Trent, 2018; Yeow *et al.*, 2018), and *in vitro* reconstitution and transport assays suggest that MlaFEDB may be bi-directional, with a preference for import (retrograde transport) (Tang *et al.*, 2020). Concurrently with our preprint, two other groups also reported structures of the MlaFEDB complex on BioRxiv (Coudray *et al.*, 2020; Mann *et al.*, 2020; Tang *et al.*, 2020), and these structures appear to be similar overall, although the PDB coordinates are not yet available to analyze. Integrating all available data, we present models for both export and import mediated by MlaFEDB, and the biggest conceptual challenge to each, which remains to be addressed in future work (**Figure 6**).

**Figure 6.**
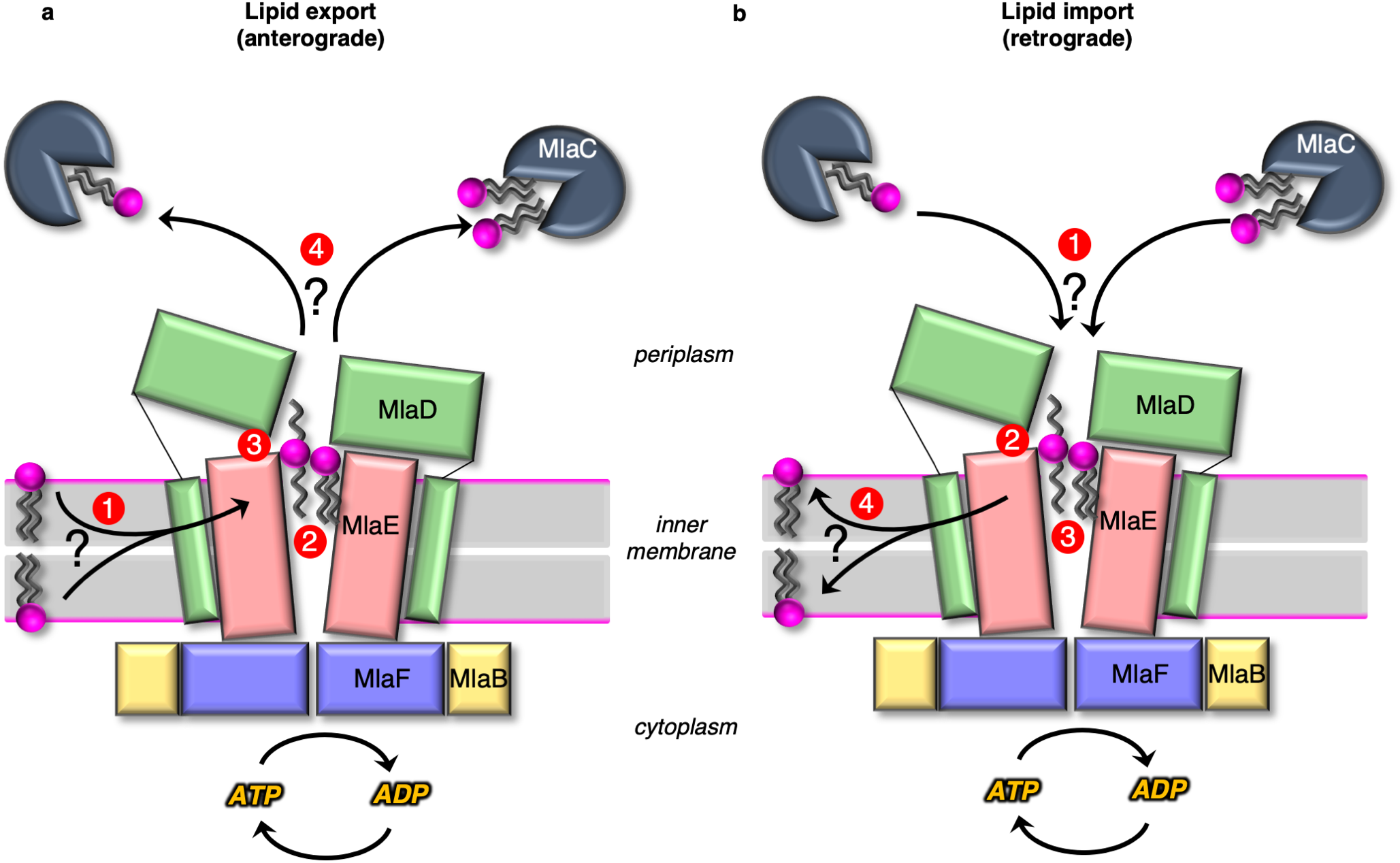
Models for lipid transport by MlaFEDB. **(a)** Lipid export model: 1) Lipids are extracted from the IM and transferred to the outward-open pocket by an unknown mechanism. 2) Lipids are reoriented, from “tails-down” to “extended” or “tails-up” configuration. 3) Conformational changes in MlaE coupled to the ATP hydrolysis cycle likely push lipids out of the MlaE pocket and into MlaD pore. 4) Lipids are transferred to MlaC to be shuttled across periplasm to the outer membrane MlaA-OmpC/F complex. MlaC may accommodate one or two phospholipids, or a single larger lipid. **(b)** Lipid import model: 1) Lipids from MlaC are transferred to MlaD, likely dependent on ATP-driven conformational changes in MlaD and MlaE. 2) Lipids travel through the continuous channel from MlaD, and are transferred to the outward-open pocket of MlaE. 3) Phospholipids are reoriented “tails-down”, as they are transported between MlaD and MlaE. 4) Lipids are inserted into the inner membrane.

In anterograde phospholipid export (**Figure 6a**), first, phospholipids must be extracted from the inner membrane and reach the outward-open pocket of MlaE, from either the inner or outer leaflet (Hughes *et al.*, 2020). Due to the structural similarities between MlaE and LptFG, which extracts LPS from the outer leaflet, it is plausible that MlaFEDB may extract PLs from the outer leaflet as well. However, it is also possible that MlaFEDB flips lipids across the bilayer from the inner leaflet to the outward-open pocket observed in our structure before exporting them. Second, substrate reorientation may need to occur such that lipids move through the MlaD pore “tails-first” to facilitate transfer from MlaD to MlaC, since MlaC binds only the lipid tails (Huang *et al.*, 2016; Ekiert *et al.*, 2017). This would require a ~180 degree flip of lipid 1 from its orientation bound to MlaE. Thus, the two lipid conformations observed in our structure may be intermediates in the process of lipid flipping, with the extended conformation of lipid 2 representing the halfway-point in the process of completely inverting the orientation of the lipid: between “tails-down” as if embedded in the outer leaflet of the IM, and “tails-up” as it traverses the MlaD pore. However, it is also possible that lipids move through the pore in an extended state (resembling lipid 2), perhaps allowing MlaC to first bind to one tail, followed by the second. The lateral channels where the head groups are located may assist in this process, and allow the accommodation of lipids with a variety of head groups. Third, by analogy to the LPS exporter (Qian *et al.*, 2017; Li, Orlando and Liao, 2019a; Owens *et al.*, 2019), we hypothesize that conformational changes in MlaE may lead to a collapse of the outward-open pocket, extruding the lipids upwards and into the MlaD pore. In the LPS exporter, this step is thought to be linked to ATP binding or hydrolysis, and it is possible that lipid reorientation and pocket collapse occur in a concerted manner, as opposed to sequentially. Fourth, the phospholipid emerges from the MlaD pore “tails-first” and is transferred to the lipid binding pocket of an MlaC protein docked on the surface of MlaD. As MlaC proteins from some species are capable of binding multiple phospholipids at one time (Ekiert *et al.*, 2017), it is possible that two lipids may be transferred from MlaFEDB to a single MlaC protein, or that multiple MlaC molecules may be involved. To complete the transport process, MlaC would then shuttle phospholipids across the periplasmic space and deliver them to the MlaA-OmpF/C complex for insertion into the outer membrane. Perhaps the biggest conceptual challenge to models for lipid export revolves around how a tightly-bound lipid can be transferred from MlaC to MlaA-OmpF/C in the absence of a direct energy source. It is unclear from available data whether this can occur spontaneously.

In retrograde phospholipid import (**Figure 6b**), first, an interaction between MlaD and MlaC must trigger the transfer of tightly-bound lipid(s) from MlaC to MlaD. Second, conformational changes in the outward-open structure of MlaE may drive the transport of lipids from the MlaD pore through the continuous channel into MlaE. Third, based on the conformations of lipid observed, it seems likely that the phospholipids will be reoriented, “tails-down” into the MlaE lipid binding site prior to being inserted into the inner membrane. Fourth, lipids must be inserted into the inner membrane. Based on the orientation in our structure, lipids in MlaE would be properly oriented for direct transfer to the outer leaflet of the inner membrane. However, the Mla system was recently reported to catalyze the flipping lipids from the outer leaflet to the inner leaflet (Hughes *et al.*, 2020), raising the possibility that imported lipids may ultimately reach the cytoplasmic leaflet. For the lipid import model, understanding the mechanism of initial lipid transfer from MlaC to MlaD is the biggest conceptual challenge. *In vitro*, lipid transfer from MlaC to MlaD has not been reported, though the reverse, lipid transfer from MlaD to MlaC, occurs spontaneously (Huang *et al.*, 2016; Ercan *et al.*, 2019; Hughes *et al.*, 2019). MlaC has a very high affinity for lipids (as evidenced by co-purification with lipids (Ercan *et al.*, 2019; Hughes *et al.*, 2019), making it likely that ATP hydrolysis is required to drive lipid transfer from MlaC to MlaD. ATP-dependent lipid transfer from MlaC would likely require coupling of ATP-hydrolysis by MlaF in the cytoplasm to the spatially distant MlaD ring. This would require coupled conformational changes transduced via the intervening MlaE subunit, to ultimately produce a conformation in MlaD that is competent to extract lipids from MlaC. Our structure does reveal conformational changes in the MlaD ring relative to the crystal structure (Ekiert *et al.*, 2017), and one possibility is that these different conformational states may be related to motions required to facilitate lipid transfer from MlaC to MlaD.

It is noteworthy that the proposed manner of lipid binding to MlaFEDB differs significantly among all three pre-prints posted around the same time. Our structure and the other *E. coli* MlaFEDB structure described by Tang, *et al* describe lipid binding to the outward-open pocket of MlaE (Tang *et al.*, 2020), while the *Acinetobacter baumannii* MlaFEDB described by Mann, *et al* describes lipid binding sites at the pore of MlaD as well as six additional lipid-binding sites in between the pore loops of each of the six MCE domains in the MlaD ring (Mann *et al.*, 2020). Tang et al observed density assigned to a single phospholipid bound in the outward-open pocket of MlaFEDB, with the head group facing towards the core of the MlaE dimer and tails pointing towards the MlaD pore. This contrasts with our observation of two lipids bound in the outward-open pocket of our structure reconstituted in lipid nanodiscs, where the lipids are bound in roughly the opposite orientation. Some of the differences between these structures could be ascribed to differences between the *E. coli* and *A. baumannii* transporters. It is also noteworthy that the sample preparation for the three structures is different. The samples for both the *A. baumannii* and Tang et al *E. coli* structures were prepared in detergent. Tang. et al also added *E. coli* lipid extract to their sample just before grid freezing. In contrast, our structure was reconstituted in lipid nanodiscs. It is unclear if these differences in lipid recognition between the two *E. coli* structures reflect differences in sample preparation, data processing methodology (e.g., asymmetric reconstruction versus application of C2 symmetry), or perhaps represent different snapshots of the transport mechanism, or even differences in lipid conformation when one vs two lipids are bound.

Our data, together with other recent studies, suggest possible mechanisms for phospholipid transport across the cell envelope, and raise the intriguing possibility that Mla may translocate multiple phospholipid substrates each transport cycle, or perhaps accommodate larger lipid substrates like cardiolipin. A defined lipid binding pocket within MlaE sets the stage for future studies of targeted inhibitors and small molecule modulators of this complex, both in the context of therapeutics against drug-resistant Gram-negative bacteria, and for the study of cell envelope biogenesis in double-membraned bacteria.

## Methods

### Expression and purification of MlaFEDB for cryo-EM

To prepare a sample for cryo-EM, plasmid pBEL1200 (Ekiert *et al.*, 2017), which contains the *mlaFEDCB* operon with an N-terminal His-tag on MlaD, was transformed into Rosetta 2 (DE3) cells (Novagen). For expression, overnight cultures of Rosetta 2 (DE3)/pBEL1200 were diluted 1:100 in LB (Difco) supplemented with carbenicillin (100 μg/mL) and chloramphenicol (38 μg/mL) and grown at 37 °C with shaking to an OD600 of 0.9, then induced by addition of L-arabinose to a final concentration of 0.2% and continued incubation for 4 hours shaking at 37 °C. Cultures were harvested by centrifugation, and the pellets were resuspended in lysis buffer (50 mM Tris pH 8.0, 300 mM NaCl, 10% glycerol). Cells were lysed by two passes through an Emulsiflex-C3 cell disruptor (Avestin), then centrifuged at 15,000 xg for 30 mins to pellet cell debris. The clarified lysates were ultracentrifuged at 37,000 rpm (F37L Fixed-Angle Rotor, Thermo-Fisher) for 45 mins to isolate membranes. The membranes were resuspended in membrane solubilization buffer (50 mM Tris pH 8.0, 300 mM NaCl, 10% glycerol, 25 mM DDM) and incubated for 1 hour, rocking at 4 °C. The solubilized membranes were then ultracentrifuged again at 37,000 rpm for 45 mins, to pellet any insoluble material. The supernatant was incubated with NiNTA resin (GE Healthcare #17531802) at 4 °C, which was subsequently washed with Ni Wash Buffer (50 mM Tris pH 8.0, 300 mM NaCl, 10 mM imidazole, 10% glycerol, 0.5 mM DDM) and bound proteins eluted with Ni Elution Buffer (50 mM Tris pH 8.0, 300 mM NaCl, 250 mM imidazole, 10% glycerol, 0.5 mM DDM). MlaFEDB containing fractions eluted from the NiNTA column were pooled and concentrated before separation on a Superdex 200 16/60 gel filtration column (GE Healthcare) equilibrated in gel filtration buffer (20 mM Tris pH 8.0, 150 mM NaCl, 0.5 mM DDM). Fractions of MlaFEDB from size exclusion chromatography were pooled and used for incorporation into nanodiscs.

### Reconstitution of MlaFEDB in lipid nanodiscs

For expression of the membrane scaffold protein, MSP1D1, overnight cultures of Rosetta 2 (DE3)/pMSP1D1 (Addgene #20061) were diluted 1:100 in LB (Difco, #244620) supplemented with kanamycin (50 μg/mL) and chloramphenicol (38 μg/mL) and grown at 37 °C with shaking to an OD600 of 0.9, then induced by addition of IPTG to a final concentration of 1 mM and continued incubation for 3 hours shaking at 37 °C. Cultures were harvested by centrifugation, and the pellets were resuspended in lysis buffer (50 mM Tris pH 8.0, 300 mM NaCl, 10 mM imidazole). Cells were lysed by two passes through an Emulsiflex-C3 cell disruptor (Avestin), then centrifuged at 38,000 xg to pellet cell debris. The clarified lysates were incubated with NiNTA resin (GE Healthcare #17531802) at 4 °C, which was subsequently washed with Ni Wash Buffer (50 mM Tris pH 8.0, 300 mM NaCl, 10 mM imidazole) and bound proteins eluted with Ni Elution Buffer (50 mM Tris pH 8.0, 300 mM NaCl, 250 mM imidazole). The His-tag was cleaved using TEV protease.

For nanodisc reconstitution, a protocol was adapted from Gao *et al(Gao et al., 2016)*. 2.5 mg of *E. coli* polar lipid extract (Avanti #100600) were dissolved in 1 ml of chloroform in a glass test tube. The chloroform was then evaporated slowly under a stream of argon gas, to produce a thin film of lipids on the bottom of the tube, and further left to dry under vacuum for at least 2 hours. The lipids were then resuspended in 200 ul of lipid resuspension buffer (20 mM HEPES, 150 mM NaCl, 14 mM DDM, pH 7.4) and sonicated until the mixture was almost clear. The lipids, MSP1D1 and MlaFEDB were mixed at a molar ratio of 400:4:1, respectively, in nanodisc buffer (20 mM HEPES, 150 mM NaCl, pH 7.4), and left to incubate on ice for 30 mins. Bio-Beads (Bio-Rad #1523920) were added (20 mg per 1 ml nanodisc mixture) and incubated for 1 hour, rocking at 4 °C. A second batch of Bio-Beads were added and incubated at 4 °C overnight. The following day, the Bio-Beads were removed and the sample separated on a Superdex 200 16/60 gel filtration column (GE Healthcare) equilibrated in nanodisc buffer (20 mM HEPES, 150 mM NaCl, pH 7.4). Fractions were assessed by SDS PAGE and negative stain EM, and were pooled and concentrated for cryo-EM grid preparation.

### Cryo-EM grid preparation and data collection

After size exclusion chromatography, 3 μL of MlaFEDB reconstituted into nanodiscs (at a final concentration of 0.95 mg/mL) was applied to 400 mesh quantifoil holey carbon grids 1.2/1.3 glow discharged for 12 seconds. The sample was then frozen in liquid ethane using the FEI Vitrobot Mark IV. Pre-screening of the grids was performed on Talos Arctica TEMs equipped with K2 cameras, operated at 200 kV, and located at PNCC (Portland, OR) or at NYU (New York, NY). Acquisition of the movies used for the final reconstruction was performed on a Titan Krios microscope (“Krios 2”) equipped with Gatan K2 Summit camera controlled via Leginon (Suloway *et al.*, 2005) and operated at 300 kV (located at the New York Structural Biology Center, NY). Image stacks of 30 frames were collected in super-resolution mode at 0.416 Å per pixel. Data collection parameters are shown in **Supplementary file 1**.

### Cryo-EM data processing

The overall strategy is summarized in supplemental **Figure 1 — figure supplement 1**. The initial preprocessing steps were all performed within Relion 2.1 (Kimanius *et al.*, 2016; Fernandez-Leiro and Scheres, 2017). Movies were drift corrected with MotionCor2 (Zheng *et al.*, 2017) and CTF estimation was performed using GCTF (Zhang, 2016). ~1,000 particles were selected manually and subjected to 2D classification. The resulting class averages were used as templates for subsequent automated particle picking of 1,283,606 particles that were extracted with a box size of 300 pixels. The data was then exported in CryoSparc v 0.6 (Punjani *et al.*, 2017) for further processing. After 2D classification, 731,205 particles were used to generate an ab-initio model subjected to heterogenous refinement of 3 classes. The 3rd class led to a map in agreement with the size and shape of a previously published low resolution reconstruction (Ekiert *et al.*, 2017). A second round of heterogeneous classification was then run with the 376,885 particles from this class: only class 3 led to a high resolution map from 209,224 particles. A curation step was applied to only include particles with assignment probability greater than a threshold of 0.95, reducing the number of particles to 177,513. Particles were then imported back to Relion for additional rounds of local refinement after having re-extracted the particles with a 500 pixel box size. In Relion 3.1-beta, we performed local CTF and aberration refinement and then performed particle polishing (re-doing first motion correction with Relion’s own implementation of MotionCorr), which improved the resolution from ~3.5 Å to ~3.3 Å. A second round of CTF and aberration refinement further improved the resolution to ~3.2 Å. The data was then imported into CryoSparc 2.12 for another round of refinement to 3.05 Å (some default parameters were modified: we used 3 extra final passes instead of 1, a batchsize epsilon of 0.0005, set the “optimize per particle defocus” and “per group ctf parameters” options to true). Average resolution was estimated using gold standard methods and implementations within Relion and Cryosparc.

Other data processing strategies were explored but failed to bring additional information or improve the resolution: signal subtraction and focused refinement of subdomains (of MlaE, or MlaFEB with and without C2 symmetry), other rounds of 3D classification, further restrict the selection of particles to the best ones relying on the probability distributions computed in Relion (rlnLogLikeliContribution rlnMaxValueProbabilityDistribution). Transfer of data from CryoSparc to Relion was performed using pyem (Asarnow, D., Palovcak, E., Cheng, Y., 2019).

### Model building

The following models were used as a starting models for the MlaFEDB structure: for MlaFB, the X-Ray structure, PDB ID: 6XGY (Kolich *et al.*, 2020); for the MCE domain protein MlaD, PDB ID: 5UW2 (Ekiert *et al.*, 2017); and for MlaE, a computationally predicted model (Ovchinnikov *et al.*, 2015; Ekiert *et al.*, 2017). Domains were docked as rigid bodies in Chimera (Pettersen *et al.*, 2004), and manual model building was done in COOT (Emsley *et al.*, 2010). The models were then iteratively refined using the real_space_refine algorithm in PHENIX (Echols *et al.*, 2012; Liebschner *et al.*, 2019), with rounds of manual model building in between using COOT. The 6 TM helices from the 6 MlaD subunits, not present in the construct used of the X-ray structure, were built *de novo*, but the loops connecting those helices to the core MCE domains were too flexible to be modelled. For the 2 MlaD TM^A/D^ helices contacting IF2 and TM3 of MlaE, as well as the 2 MlaD TM^B/E^ helices that contact IF1 on MlaE, the clear side chain density allowed unambiguous assignment of the sequence register. For the final MlaD TM^C/F^ helices contacting IF1 of MlaE near the N-Terminus, the EM map was filtered to 6 Å to better visualize the density, as these helices are more flexible. The close helical packing geometry between IF1 and MlaD-TM^B/E^ enforces a strong preference for glycine residues at the positions of closest contact (Gly21^MlaE^ and Gly11^MlaD^, ~3.4 Å C -C) (**Figure 3 — figure supplement 1a**). Gly is present at MlaD position 11 in 13/13 sequences analyzed, and at MlaE position 21 in 12/13 sequences analyzed (**Figure 3 — figure supplement 1b, c**), suggesting that the interactions between IF1 and MlaD-TM^B/E^ are specific and conserved. Gly11 of MlaD-TM^B/E^ and residues of IF1 are part of a larger interaction motif (17-LxxFGxxxL-25) (**Figure 3 — figure supplement 1a**). MlaD-TM^C/F^ appears to interact with IF1 in a similar manner in the vicinity of MlaE Gly10 (Gly10^MlaE^ and Gly11^MlaD^, ~3.5 Å C -C), which is part of a very similar conserved motif (6-LxxLGxxxI-14; **Figure 3 — figure supplement 1**). While density for side chains in MlaD TM^C/F^ is weak, the similarity in helix packing geometry and these two binding sites, along with only one available Gly for close helix packing in the MlaD TM helix suggest that the same surface of the MlaD TM is used for IF1 binding in these chains C and F as well. Consequently, we have used this Gly-Gly close packing to establish a likely sequence register for these TM helices. Due to the lower resolution, we did not model the side chains of these residues explicitly.

The MlaE region displayed the highest local resolution (below 3 Å in its core) and was almost entirely modelled. The two most flexible regions were the extremities of the interfacial helices IF1 (the N-Terminus and the connection to the IF2/TM1 helix). Due to a lack of density on the N-terminus of IF1, residues 1-4 could not be modeled. Within the MlaE dimer and at the interface with MlaD, we identified two clearly defined densities that corresponded to the shape and size of phospholipids, which are present in our reconstitution. Two phosphatidylethanolamine molecules (code: PEF) were manually placed into the densities. The phosphate, glycerol backbone, and most of the C16 fatty acid chains could readily be placed in the map, but the ethanolamine portion of the head group was removed due to a lack of density, meaning this part may be flexible and/or non-specific to a certain type of lipid. In reality, our MlaFEDB sample likely contained a heterogeneous mixture of lipids bound, with a range of head groups and acyl chain lengths/unsaturations. Although the resulting ligands resemble phosphatidic acid, we retain the PEF/phosphatidylethanolamine designation, as PE is the most abundant PL species in *E. coli* but phosphatidic acid is relatively scarce.

We have modeled two PE molecules bound to the transporter simultaneously, as this best explains the all the available information, including: 1) The two densities are well-resolved and do not cross each other; 2) after refinement, the atoms of the lipids are roughly within van der Waals distance of each other and nearby protein atoms, without excessive clashes and in line with expectations for flexible/heterogeneous ligands at this resolution; 3) while the tight packing of two lipids fills the MlaE pocket, binding of a single lipid would the pocket ~50% empty; based upon the observed protein-lipid interactions, it is difficult to envision how single lipid molecules could be bound in the pocket yet be constraint of the observed conformation of lipid 1 and lipid 2 unless a second lipid is simultaneously present; 4) we performed various focused 3D classification with variable masks and regularization parameters in Relion, as well as 3D variability analysis in cluster mode in Cryosparc with and without a mask. While the resulting maps were generally of lower quality, the reconstructions containing clear lipid-like densities most closely resembled the configuration of the two lipids modeled in our structure.

Using both the high resolution map and its filtered version at 6 Å, we also modelled both coils of the membrane scaffold protein belt surrounding the edges of the nanodisc (starting with the ones modelled in PDB: 6CM1). These MSP belts took the form of two relatively featureless tubes of density. Consequently, their position is modeled as using polyalanine helices, and we were able to account for ~160 of the expected ~190 residues for MSP1D1.

The final model of the MlaFEDB complex is nearly complete, with two noteworthy areas of disorder. First, the loops between the TM helices and the MCE domains of MlaD could not be resolved (5-8 residues disordered in each). Second, the last 32 residues at the C-terminus of each MlaD chain (residues 153-183), which were disordered in previous X-ray structures (Ekiert *et al.*, 2017), were also not visible in our EM structure.

### Structure analysis and Bioinformatics

As structural deviations between MlaE and other ABC TMDs made database searches more difficult, we conducted a Dali search (Holm, 2019) initiated with MlaF to recover all of the PDB structures containing an ABC domain. These structures were then manually curated based upon the presence or absence of TMDs and further classified based upon the TMD fold.

In order to assess the conservation of MlaD and MlaE sequences across species, we identified at least one “representative” species from each major bacterial order, across the entire bacterial kingdom. Within each order, we selected “representative” species, which was typically one of the most widely studied members and/or a special most impacting human health (prior to an examination of the sequences, to avoid bias). For each representative species, we searched the reference genome using BLAST to identify possible homologs of *E. coli* MlaE, MlaD, MlaC, and MlaA. We did not search directly for MlaF or MlaB homologs, as ABC ATPases and STAS domain proteins unrelated to Mla are common in bacteria. Of 65 species analyzed, only 13 were determined to encode what appeared to be functional MCE transporters that were “true homologs” of Mla (**Supplementary file 3)**. To be included in this group, the species must encode a homolog of MlaD (single MCE domain without a long C-terminal helical region (less than ~50 residues) and also homologs of MlaE, MlaC, and MlaA in its genome. In every case, MlaE, MlaD, and MlaC were encoded just downstream of an MlaF-like ABC subunit, and just upstream of an MlaB-like protein (except in *Rickettsia rickettsii*, which appears to lack MlaB). Sometimes MlaA was encoded in the same operon, while sometimes it was encoded elsewhere in the genome. The resulting “True Mla” homologs were subsequently used for sequence analysis. Sequence alignments were generated using MUSCLE (Edgar, 2004) and visualized using JalView (Waterhouse *et al.*, 2009).

The 3DFSC in **Supplementary file 2** was measured using the Remote 3DFSC Processing Server (Tan *et al.*, 2017). The interfaces between the different subunits of MlaFEDB, Lpt and ABCA/G proteins were analyzed using the COCOMAPS server (Vangone *et al.*, 2011). The area of the cavities of MlaE and LptFG were estimated using CASTp (Tian *et al.*, 2018) and HOLLOW (Ho and Gruswitz, 2008). The curvature of the IF2-TM1 helices was analyzed using Bendix (Dahl, Chavent and Sansom, 2012) and the corresponding figures generated with VMD software support which is developed with NIH support by the Theoretical and Computational Biophysics group at the Beckman Institute, University of Illinois at Urbana-Champaign. All other figures were made with Chimera (Pettersen *et al.*, 2004) or PyMOL (Schrödinger, LLC). The PyMOL plugin, anglebetweenhelices (Schrödinger, LLC), was used to compute the angle between IF1 of MlaE and the TM helices of MlaD.

### Phenotypic assays for *mla* mutants in *E. coli*

Knockouts of *mlaD* and *mlaE* were constructed in *E. coli* BW25113 by P1 transduction from the Keio collection (Baba *et al.*, 2006), followed by excision of the antibiotic resistance cassettes using pCP20 (Cherepanov and Wackernagel, 1995). Serial dilutions of the strains in 96 well plates were manually spotted (2 uL each) on plates containing LB agar or LB agar supplemented with 0.2% SDS and 0.35 mM EDTA, and incubated for 16 hours at 37 °C. We find that this growth assay is very sensitive to the reagents used, particularly the LB agar (see Kolich *et al (Kolich et al., 2020)*). For the experiments reported here, we used Difco LB agar pre-mix (BD Difco #244510), a 10% stock solution of SDS (Sigma L5750), and a 500 mM stock solution of EDTA, pH 8.0 (Sigma ED2SS). Furthermore, we note that when the agar plates were incubated longer than 16 hours, we began to observe some clearing/loss of pigmentation of the bacterial spot dilutions.

For complementation and/or testing the functionality of the various MlaD and MlaE mutants, a pBAD-derived plasmid harboring the *mlaFEDCB* operon was transformed into the appropriate knockout strain. To test the functionality of mutations in MlaD, we transformed the *mlaD* knockout strain with pBEL1198 (*mlaFEDCB* operon N-terminal His-tag on MlaE, see **Supplementary file 4**), or derivatives of this plasmid harboring the desired mutations in MlaD (MlaD TM replaced with LptC TM (pBEL2139) and MlaD TM replaced with LetB TM (pBEL2138), see **Supplementary file 4)** For the MlaE mutants, we transformed the *mlaE* knockout strain with pBEL1200 (*mlaFEDCB* operon with N-terminal His-tag on MlaD, see **Supplementary file 4**), or derivatives of this plasmid harboring the desired mutations in MlaE (IF1 1-15 aa deletion (pBEL2093), IF1 1-25 aa deletion (pBEL2132), IF1 1-30 aa deletion (pBEL2092), IF1 1-39 aa deletion (pBEL2133), Tyr81Ala (pBEL2099), Tyr81Trp (pBEL2100), Arg97Ala (pBEL2098), Glu98Ala (pBEL2094), Lys205Ala (pBEL2095) and Asp250Ala (pBEL2097), see **Supplementary file 4)**. We found that leaky expression from the pBAD promoter was sufficient for complementation of the phenotypes of both the *mlaD* and *mlaE* knockout strains, and thus no L-arabinose was added. We suspect that these constructs significantly over-produce MlaFEDCB proteins, and while some mutants tested confirmed our ability to detect loss-of-function mutations, it is possible that this over-expression may mask the impact of mutations that cause a moderate reduction in MlaFEDB activity.

### Expression and purification of MlaFEDB mutants

MlaFEDB mutants were expressed, and purified by NiNTA affinity chromatography as described above. For studies involving mutations in MlaD, we used a construct with the WT *mlaFEDCB* operon with an N-terminal His-tag on MlaE (pBEL1198), or derivatives in which the MlaD TM was replaced with LptC TM (pBEL2139) or LetB TM (pBEL2138). For studies involving mutations in MlaE, we used a construct with the WT *mlaFEDCB* operon with an N-terminal His-tag on MlaD (pBEL1200), or derivatives with the MlaE IF1 1-15 aa deletion (pBEL2093), MlaE IF1 1-25 aa deletion (pBEL2132), MlaE IF1 1-30 aa deletion (pBEL2092), or MlaE IF1 1-39 aa deletion (pBEL2133).

### Western blot to detect MlaD TM mutants

In order to assess the expression and cellular localisation of the MlaD mutants with the native TM replaced by the TM from LetB or LptC, each of the pBEL1198 derived plasmids (WT operon (pBEL1198), MlaD TM replaced with LptC TM (pBEL2139) and MlaD TM replaced with LetB TM (pBEL2138), see **Supplementary file 4)** were expressed and purified as described above (see **Expression and purification of MlaFEDB**). Following cell lysis and a low speed spin to remove cell debris, a sample was collected, which we refer to as the “whole cell lysate”. The membranes were then isolated and solubilized as described above, and a sample was taken from the solubilized membranes, which we refer to as the “membrane fraction”. 10 μl of each sample were separated on an SDS-PAGE gel and transferred to a nitrocellulose membrane. The membranes were blocked in PBST containing 5% milk for 1 hour. The membranes were then incubated with primary antibody (rabbit polyclonal anti-MlaD (provided by Henderson lab, University of Queensland) at a dilution of 1:10000) in PBST + 5% milk overnight at 4 °C. The membranes were then washed 3 times with PBST and were incubated with goat anti-rabbit IgG polyclonal antibody (IRDye^®^ 800CW, LI-COR Biosciences #925-32211 at a dilution of 1:10000) in PBST + 5% milk for 1 hour. The membranes were then washed 3 times with PBST and imaged using a LI-COR (LI-COR Biosciences).

### Lipid crosslinking experiments

This method was adapted from Isom and Coudray *et al*, 2020 (Isom *et al.*, 2020). T7express *E. coli* (NEB) were co-transformed with 1) plasmids to express MlaFEDCB (either the WT proteins using pBEL1198, or derivatives of this plasmid expressing Amber mutant MlaE variants for BPA incorporation (Tyr81Bpa (pBEL2057), Val77Bpa (pBEL2060), Leu78Bpa (pBEL2061), Leu70Bpa (pBEL2062), Leu99Bpa (pBEL2063), Trp149Bpa (pBEL2065) or Phe209Bpa (pBEL2066)); and 2) pEVOL-pBpF (Addgene #31190), which encodes a tRNA synthetase/tRNA pair for the *in vivo* incorporation p-benzoyl-l-phenylalanine (BPA) in *E. coli* proteins at the amber stop codon, TAG (Chin *et al.*, 2002; Isom *et al.*, 2017). Bacterial colonies were inoculated in LB broth supplemented with carbenicillin (100 μg/mL), chloramphenicol (38 μg/mL) and 1% glucose, and grown overnight at 37 °C. The following day, bacteria were pelleted and resuspended in ^32^P Labelling Medium (a low phosphate minimal media: 1 mM Na2HP04, 1 mM KH2PO4, 50 mM NH4Cl, 5 mM Na2SO4, 2 mM MgSO4, 20 mM Na2-Succinate, 0.2x trace metals and 0.2% glucose) supplemented with carbenicillin (100 μg/mL) and chloramphenicol (38 μg/mL) and inoculated 1:33 in the 10 ml of the same medium. Bacteria were grown to OD 1.0 and a final concentration of 0.2% L-arabinose and 0.5 mM BPA (Bachem, #F-2800.0005), alongside 375 μCi ^32^P orthophosphoric acid (PerkinElmer, #NEX053010MC) were added and left to induce overnight.

The following day, the cultures were spun down and resuspended in 1ml of PBS, and the “crosslinked” samples underwent crosslinking by treatment with 365 nM UV in a Spectrolinker for 30 mins. Both the crosslinked and uncrosslinked cells were then spun down and resuspended in 1 ml of lysozyme-EDTA resuspension buffer (50 mM Tris pH 8.0, 300 mM NaCl, 10 mM imidazole, 1 mg/ml lysozyme, 0.5 mM EDTA, 25U/ml benzonase) and were incubated for 1 hour at room temperature. The cells then underwent 8 cycles of freeze-thaw lysis by alternating between liquid nitrogen and a 37 °C heat block. The lysate was pelleted at 20,000 xg for 15 minutes, and the pellets were resuspended in 133 μl of membrane resuspension buffer (50 mM Tris pH 8.0, 300 mM NaCl, 10% glycerol and 25 mM DDM), and incubated, shaking, for 1 hour. The sample volume was then increased to 1 ml using 10 mM wash buffer (50 mM Tris pH 8.0, 300 mM NaCl, 10 mM imidazole) and insoluble material was pelleted at 20,000 xg for 15 minutes. Each supernatant was then mixed with 25 μl of nickel beads (Ni Sepharose 6 Fast Flow) for 30 mins. The beads were pelleted at 500 xg for 1 min and the supernatant collected. The beads were then washed four times with 40 mM wash buffer (50 mM Tris pH 8.0, 300 mM NaCl, 40 mM imidazole, 10% glycerol, 0.5 mM DDM) and finally resuspended in 50 μl of elution buffer (50 mM Tris pH 8.0, 300 mM NaCl, 300 mM imidazole, 10% glycerol, 0.5 mM DDM). The samples were then mixed with 5x SDS-PAGE loading buffer, and the beads spun down at 12,000 xg for 2 mins. Eluted protein was analyzed by SDS-PAGE and stained using InstantBlue™ Protein Stain (Expedeon, #isb1l). Relative loading of the MlaE monomer band on the gel was estimated integrating the density of the corresponding bands in the InstantBlue-stained gel in ImageJ (Rueden *et al.*, 2017), and this was used to normalize the amount of MlaE monomer loaded on a second gel, to enable more accurate comparisons between samples. The normalized gel was stained with InstantBlue and ^32^P signal was detected using a phosphor screen and scanned on a Typhoon scanner (Amersham). Three replicates of the experiment were performed, starting with protein expression. NB: earlier protocols using urea solubilization (Coudray *et al.*, 2020) gave globally similar results but with variation in cross linking efficiency between biological replicates; the improved protocol used here, purifying MlaFEDB under native conditions (without urea), has much lower variation between replicates.

### Western blots to detect MlaE in crosslinked samples

Samples were grown and induced as described above (see **Lipid crosslinking experiments**), but in the absence of ^32^P orthophosphoric acid. The following day, the cultures were spun down and resuspended in 500 μl of lysozyme-EDTA resuspension buffer (50 mM Tris pH 8.0, 300 mM NaCl, 10 mM imidazole, 1 mg/ml lysozyme, 0.5 mM EDTA, 25U benzonase) and were incubated for 1 hour at room temperature. The samples then underwent crosslinking by treatment with 365 nM UV in a Spectrolinker for 30 mins. For lysis, the crosslinked samples were added to 250 μl of 100 mM DDM, and 0.48 g of urea, and adjusted up to a total volume of 1 ml using 10 mM wash buffer (50 mM Tris pH 8.0, 300 mM NaCl, 10 mM imidazole), and incubated at 60 °C, with intermittent inversion of the tubes to mix, until the urea was dissolved and the cells had undergone lysis (approximately 15 mins). Each sample was then mixed with 25 μl of NiNTA resin (Ni Sepharose 6 Fast Flow) for 30 mins. The resin was pelleted at 500 xg for 1 min and the supernatant collected. The resin was then washed four times with urea wash buffer (50 mM Tris pH 8.0, 300 mM NaCl, 40 mM imidazole, 8 M urea, 0.5 mM DDM) and finally resuspended in 50 μl of urea wash buffer (50 mM Tris pH 8.0, 300 mM NaCl, 250 mM imidazole, 8 M urea, 0.5 mM DDM). The samples were mixed with 5x SDS-PAGE loading buffer, and the resin spun down at 12,000 xg for 2 mins. The Western blot was done as described above (in **Western blots to detect MlaD TM mutants**), but with an anti-His antibody (Qiagen, #34660 at a dilution of 1:5000) as primary, to detect His-tagged MlaE, and HRP-linked anti-mouse (GE Healthcare, #NA931-1ML, at a dilution of 1:5000) as the secondary antibody, and was developed using a Bio-Rad ChemiDoc imager.

## Acknowledgments

We thank Marisa Lopez-Redondo (NYU) and Yuan Gao (UCSF) for guidance with nanodisc reconstitution, and members of the Bhabha/Ekiert labs for helpful discussions. We thank Ian Henderson (University of Queensland) for providing anti-MlaD antibodies. We thank Noelle Antao, Juliana Ilmain, Dhenesh Puvanendren, Rachel Redler and Casey Vieni for critical reading and feedback on our manuscript. We gratefully acknowledge the following funding sources: NIH grant R35GM128777 (D.C.E.), NIH grant R00GM112982, Damon Runyon Cancer Research Foundation grant DFS-20-16 and Pew Charitable Trusts PEW-00033055 (G.B.). American Heart Association postdoctoral fellowship 20POST35210202 (G.L.I), NIH T32 predoctoral training grant T32 GM088118 (M.R.M). Pre-screening for cryoEM sample optimization was carried out at the NYU cryo-EM core facility and the Pacific Northwest Center for Cryo EM. CryoEM data were collected at the Simons Electron Microscopy Center and National Resource for Automated Molecular Microscopy located at the New York Structural Biology Center, supported by grants from the Simons Foundation (SF349247), NYSTAR, and the NIH National Institute of General Medical Sciences (GM103310) with additional support from Agouron Institute (F00316) and NIH (OD019994). A portion of this research was supported by NIH grant U24GM129547 and performed at the Pacific Northwest Center for Cryo-EM at Oregon Health & Sciences University (OHSU) and accessed through EMSL (grid.436923.9), a DOE Office of Science User Facility sponsored by the Office of Biological and Environmental Research. EM data processing has utilized computing resources at the HPC Facility at NYU, and we thank the HPC team. Molecular graphics and analyses performed with UCSF Chimera, developed by the Resource for Biocomputing, Visualization, and Informatics at the University of California, San Francisco, with support from NIH P41-GM103311.

## Author contributions

Conceptualization: G.L.I., M.R.M., N.C., G.B. and D.C.E; Validation: G.L.I., M.R.M., N.C., G.B. and D.C.E; Formal Analysis: G.L.I., M.R.M., N.C., G.B. and D.C.E; Investigation: G.L.I., M.R.M., N.C., G.B., M.N.S.; Writing – Original Draft Preparation: G.L.I, M.R.M., N.C., G.B. and D.C.E; Writing – Review & Editing: G.L.I., M.R.M., N.C., M.N.S., G.B. and D.C.E; Visualization: N.C.; Supervision: G.B. and D.C.E.; Funding Acquisition: G.L.I., G.B. and D.C.E,.

## Supplementary file legends

**Supplementary file 1. Data collection parameters for cryoEM structure of MlaFEDB**

**Supplementary file 2. Data refinement statistics for cryo EM structure of MlaFEDB**

**Supplementary file 3. Species included in analysis of Mla sequence conservation**

**Supplementary file 4. Plasmids used in this study**

## References

Abellón-Ruiz, J. et al. (2017) ‘Structural basis for maintenance of bacterial outer membrane lipid asymmetry’, Nature microbiology, 2(12), pp. 1616–1623.

Aller, S. G. et al. (2009) ‘Structure of P-glycoprotein reveals a molecular basis for poly-specific drug binding’, Science, 323(5922), pp. 1718–1722.

Asarnow, D., Palovcak, E., Cheng, Y. (2019) UCSF pyem, UCSF pyem v0.5. doi: 10.5281/zenodo.3576630.

Baba, T. et al. (2006) ‘Construction of Escherichia coli K-12 in-frame, single-gene knockout mutants: the Keio collection’, Molecular systems biology, 2, p. 2006.0008.

Bayburt, T. H., Grinkova, Y. V. and Sligar, S. G. (2002) ‘Self-Assembly of Discoidal Phospholipid Bilayer Nanoparticles with Membrane Scaffold Proteins’, Nano letters. American Chemical Society, 2(8), pp. 853–856.

Bi, Y. et al. (2018) ‘Architecture of a channel-forming O-antigen polysaccharide ABC transporter’, Nature, 553(7688), pp. 361–365.

Caffalette, C. A. et al. (2019) ‘A lipid gating mechanism for the channel-forming O antigen ABC transporter’, Nature communications, 10(1), p. 824.

Chen, L. et al. (2020) ‘Cryo-electron Microscopy Structure and Transport Mechanism of a Wall Teichoic Acid ABC Transporter’, mBio. doi: 10.1128/mbio.02749-19.

Cherepanov, P. P. and Wackernagel, W. (1995) ‘Gene disruption in Escherichia coli: TcR and KmR cassettes with the option of Flp-catalyzed excision of the antibiotic-resistance determinant’, Gene, pp. 9–14. doi: 10.1016/0378-1119(95)00193-a.

Chin, J. W. et al. (2002) ‘Addition of a photocrosslinking amino acid to the genetic code of Escherichia coli’, Proceedings of the National Academy of Sciences, pp. 11020–11024. doi: 10.1073/pnas.172226299.

Chong, Z.-S., Woo, W.-F. and Chng, S.-S. (2015) ‘Osmoporin OmpC forms a complex with MlaA to maintain outer membrane lipid asymmetry in Escherichia coli’, Molecular microbiology, 98(6), pp. 1133–1146.

Coudray, N. et al. (2020) ‘Structure of MlaFEDB lipid transporter reveals an ABC exporter fold and two bound phospholipids’, bioRxiv. doi: 10.1101/2020.06.02.129247.

Crow, A. et al. (2017) ‘Structure and mechanotransmission mechanism of the MacB ABC transporter superfamily’, Proceedings of the National Academy of Sciences of the United States of America, 114(47), pp. 12572–12577.

Dahl, A. C. E., Chavent, M. and Sansom, M. S. P. (2012) ‘Bendix: intuitive helix geometry analysis and abstraction’, Bioinformatics, 28(16), pp. 2193–2194.

Echols, N. et al. (2012) ‘Graphical tools for macromolecular crystallography in PHENIX’, Journal of applied crystallography, 45(Pt 3), pp. 581–586.

Edgar, R. C. (2004) ‘MUSCLE: multiple sequence alignment with high accuracy and high throughput’, Nucleic Acids Research, pp. 1792–1797. doi: 10.1093/nar/gkh340.

Ekiert, D. C. et al. (2017) ‘Architectures of Lipid Transport Systems for the Bacterial Outer Membrane’, Cell, 169(2), pp. 273–285.e17.

Emsley, P. et al. (2010) ‘Features and development of Coot’, Acta crystallographica. Section D, Biological crystallography, 66(Pt 4), pp. 486–501.

Ercan, B. et al. (2019) ‘Characterization of Interactions and Phospholipid Transfer between Substrate Binding Proteins of the OmpC-Mla System’, Biochemistry, 58(2), pp. 114–119.

Fernandez-Leiro, R. and Scheres, S. H. W. (2017) ‘A pipeline approach to single-particle processing in RELION’, Acta crystallographica. Section D, Structural biology, 73(Pt 6), pp. 496–502.

Fitzpatrick, A. W. P. et al. (2017) ‘Structure of the MacAB-TolC ABC-type tripartite multidrug efflux pump’, Nature microbiology, 2, p. 17070.

Gao, Y. et al. (2016) ‘TRPV1 structures in nanodiscs reveal mechanisms of ligand and lipid action’, Nature, 534(7607), pp. 347–351.

Gerber, S. et al. (2008) ‘Structural basis of trans-inhibition in a molybdate/tungstate ABC transporter’, Science, 321(5886), pp. 246–250.

Gu, Y. et al. (2016) ‘Structural basis of outer membrane protein insertion by the BAM complex’, Nature, 531(7592), pp. 64–69.

Han, L. et al. (2016) ‘Structure of the BAM complex and its implications for biogenesis of outer-membrane proteins’, Nature structural & molecular biology, 23(3), pp. 192–196.

Ho, B. K. and Gruswitz, F. (2008) ‘HOLLOW: generating accurate representations of channel and interior surfaces in molecular structures’, BMC structural biology, 8, p. 49.

Holm, L. (2019) ‘Benchmarking fold detection by DaliLite v.5’, Bioinformatics, 35(24), pp. 5326–5327.

Huang, Y.-M. M. et al. (2016) ‘Molecular dynamic study of MlaC protein in Gram-negative bacteria: conformational flexibility, solvent effect and protein-phospholipid binding’, Protein Science, pp. 1430–1437. doi: 10.1002/pro.2939.

Hughes, G. W. et al. (2019) ‘Evidence for phospholipid export from the bacterial inner membrane by the Mla ABC transport system’, Nature microbiology, 4(10), pp. 1692–1705.

Hughes, G. W. et al. (2020) ‘MlaFEDB displays flippase activity to promote phospholipid transport towards the outer membrane of Gram-negative bacteria’, bioRxiv. doi: 10.1101/2020.06.06.138008.

Isom, G. L. et al. (2017) ‘MCE domain proteins: conserved inner membrane lipid-binding proteins required for outer membrane homeostasis’, Scientific reports, 7(1), p. 8608.

Isom, G. L. et al. (2020) ‘LetB Structure Reveals a Tunnel for Lipid Transport across the Bacterial Envelope’, Cell, 181(3), pp. 653–664.e19.

Kadaba, N. S. et al. (2008) ‘The high-affinity E. coli methionine ABC transporter: structure and allosteric regulation’, Science, 321(5886), pp. 250–253.

Kamischke, C. et al. (2019) ‘The Acinetobacter baumannii Mla system and glycerophospholipid transport to the outer membrane’, eLife. doi: 10.7554/elife.40171.

Karpowich, N. et al. (2001) ‘Crystal structures of the MJ1267 ATP binding cassette reveal an induced-fit effect at the ATPase active site of an ABC transporter’, Structure, 9(7), pp. 571–586.

Kimanius, D. et al. (2016) ‘Accelerated cryo-EM structure determination with parallelisation using GPUs in RELION-2’, eLife, 5. doi: 10.7554/eLife.18722.

Knowles, T. J. et al. (2009) ‘Membrane protein architects: the role of the BAM complex in outer membrane protein assembly’, Nature reviews. Microbiology, 7(3), pp. 206–214.

Kolich, L. et al. (2020) ‘Structure of MlaFB uncovers novel mechanisms of ABC transporter regulation’, bioRxiv. doi: 10.1101/2020.04.27.064196.

Lee, J.-Y. et al. (2016) ‘Crystal structure of the human sterol transporter ABCG5/ABCG8’, Nature, 533(7604), pp. 561–564.

Liebschner, D. et al. (2019) ‘Macromolecular structure determination using X-rays, neutrons and electrons: recent developments in Phenix’, Acta crystallographica. Section D, Structural biology, 75(Pt 10), pp. 861–877.

Li, Y., Orlando, B. J. and Liao, M. (2019a) ‘Structural basis of lipopolysaccharide extraction by the LptB2FGC complex’, Nature, pp. 486–490. doi: 10.1038/s41586-019-1025-6.

Li, Y., Orlando, B. J. and Liao, M. (2019b) ‘Structural basis of lipopolysaccharide extraction by the LptBFGC complex’, Nature, 567(7749), pp. 486–490.

Malinverni, J. C. and Silhavy, T. J. (2009) ‘An ABC transport system that maintains lipid asymmetry in the gram-negative outer membrane’, Proceedings of the National Academy of Sciences of the United States of America, 106(19), pp. 8009–8014.

Mann, D. et al. (2020) ‘Structural basis for lipid transport by the MLA complex’, bioRxiv. doi: 10.1101/2020.05.30.125013.

Manolaridis, I. et al. (2018) ‘Cryo-EM structures of a human ABCG2 mutant trapped in ATP-bound and substrate-bound states’, Nature, 563(7731), pp. 426–430.

Murakami, S., Okada, U. and Yamashita, E. (2017) ‘Crystal structure of tripartite-type ABC transporter, MacB from Acinetobacter baumannii’. doi: 10.2210/pdb5gko/pdb.

Neumann, J., Rose-Sperling, D. and Hellmich, U. A. (2017) ‘Diverse relations between ABC transporters and lipids: An overview’, Biochimica et Biophysica Acta, Biomembranes, 1859(4), pp. 605–618.

Okuda, S. et al. (2016) ‘Lipopolysaccharide transport and assembly at the outer membrane: the PEZ model’, Nature reviews. Microbiology, 14(6), pp. 337–345.

Okuda, S., Freinkman, E. and Kahne, D. (2012) ‘Cytoplasmic ATP Hydrolysis Powers Transport of Lipopolysaccharide Across the Periplasm in E. coli’, Science, pp. 1214–1217. doi: 10.1126/science.1228984.

Orelle, C. et al. (2010) ‘Dynamics of α-helical subdomain rotation in the intact maltose ATP-binding cassette transporter’, Proceedings of the National Academy of Sciences of the United States of America. National Academy of Sciences, 107(47), pp. 20293–20298.

Ovchinnikov, S. et al. (2015) ‘Large-scale determination of previously unsolved protein structures using evolutionary information’, eLife, 4, p. e09248.

Owens, T. W. et al. (2019) ‘Structural basis of unidirectional export of lipopolysaccharide to the cell surface’, Nature, 567(7749), pp. 550–553.

Pettersen, E. F. et al. (2004) ‘UCSF Chimera--a visualization system for exploratory research and analysis’, Journal of computational chemistry, 25(13), pp. 1605–1612.

Powers, M. J. and Trent, M. S. (2018) ‘Phospholipid retention in the absence of asymmetry strengthens the outer membrane permeability barrier to last-resort antibiotics’, Proceedings of the National Academy of Sciences of the United States of America, 115(36), pp. E8518–E8527.

Punjani, A. et al. (2017) ‘cryoSPARC: algorithms for rapid unsupervised cryo-EM structure determination’, Nature methods, 14(3), pp. 290–296.

Qian, H. et al. (2017) ‘Structure of the Human Lipid Exporter ABCA1’, Cell, 169(7), pp. 1228–1239.e10.

Rueden, C. T. et al. (2017) ‘ImageJ2: ImageJ for the next generation of scientific image data’, BMC bioinformatics, 18(1), p. 529.

Shrivastava, R. and Chng, S.-S. (2019) ‘Lipid trafficking across the Gram-negative cell envelope’, Journal of Biological Chemistry, pp. 14175–14184. doi: 10.1074/jbc.aw119.008139.

Shrivastava, R., Jiang, X. ’er and Chng, S.-S. (2017) ‘Outer membrane lipid homeostasis via retrograde phospholipid transport in Escherichia coli’, Molecular microbiology, 106(3), pp. 395–408.

Smith, P. C. et al. (2002) ‘ATP binding to the motor domain from an ABC transporter drives formation of a nucleotide sandwich dimer’, Molecular cell, 10(1), pp. 139–149.

Sperandeo, P., Martorana, A. M. and Polissi, A. (2017) ‘Lipopolysaccharide biogenesis and transport at the outer membrane of Gram-negative bacteria’, Biochimica et Biophysica Acta (BBA) – Molecular and Cell Biology of Lipids, pp. 1451–1460. doi: 10.1016/j.bbalip.2016.10.006.

Suloway, C. et al. (2005) ‘Automated molecular microscopy: the new Leginon system’, Journal of structural biology, 151(1), pp. 41–60.

Tang, X. et al. (2019) ‘Cryo-EM structures of lipopolysaccharide transporter LptB2FGC in lipopolysaccharide or AMP-PNP-bound states reveal its transport mechanism’, Nature Communications. doi: 10.1038/s41467-019-11977-1.

Tang, X. et al. (2020) ‘Structural insight into outer membrane asymmetry maintenance of Gram-negative bacteria by the phospholipid transporter MlaFEDB’, bioRxiv. doi: 10.1101/2020.06.04.133611.

Tan, Y. Z. et al. (2017) ‘Addressing preferred specimen orientation in single-particle cryo-EM through tilting’, Nature methods, 14(8), pp. 793–796.

Thomas, C. et al. (2020) ‘Structural and functional diversity calls for a new classification of ABC transporters’, FEBS letters.

Thong, S. et al. (2016) ‘Defining key roles for auxiliary proteins in an ABC transporter that maintains bacterial outer membrane lipid asymmetry’, eLife, 5. doi: 10.7554/eLife.19042.

Tian, W. et al. (2018) ‘CASTp 3.0: computed atlas of surface topography of proteins’, Nucleic acids research, 46(W1), pp. W363–W367.

Vangone, A. et al. (2011) ‘COCOMAPS: a web application to analyze and visualize contacts at the interface of biomolecular complexes’, Bioinformatics, 27(20), pp. 2915–2916.

Waterhouse, A. M. et al. (2009) ‘Jalview Version 2--a multiple sequence alignment editor and analysis workbench’, Bioinformatics, pp. 1189–1191. doi: 10.1093/bioinformatics/btp033.

Yeow, J. et al. (2018) ‘The architecture of the OmpC– MlaA complex sheds light on the maintenance of outer membrane lipid asymmetry inEscherichia coli’, Journal of Biological Chemistry, pp. 11325–11340. doi: 10.1074/jbc.ra118.002441.

Zhang, K. (2016) ‘Gctf: Real-time CTF determination and correction’, Journal of Structural Biology, pp. 1–12. doi: 10.1016/j.jsb.2015.11.003.

Zheng, S. Q. et al. (2017) ‘MotionCor2: anisotropic correction of beam-induced motion for improved cryo-electron microscopy’, Nature Methods, pp. 331–332. doi: 10.1038/nmeth.4193.

