## Supplemental file 4 for "Structure of bacterial phospholipid transporter MlaFEDB with substrate bound"

**Supplementary file 4: Plasmids used in this study**

| Plasmid ID | Use of plasmid | Source |
| --- | --- | --- |
| pBEL1200  (Addgene ID: 138526) | MlaF-MlaE-(6xHis-MlaD)-MlaC-MlaB: Expression and complementation plasmid for MlaFEDCB, with N-terminal his tag on MlaD | Ekiert *et al*, 2017 |
| pMSP1D1  (Addgene ID: 20061) | Expression construct from membrane scaffold protein MSP1D1 | Denisov *et al.*, 2004 |
| pCP20 | Plasmid for removal of kanamycin resistance cassette | [Cherepanov](https://pubmed.ncbi.nlm.nih.gov/?term=Cherepanov+PP&cauthor_id=7789817) *et al.*, 1995 |
| pBEL2139 | MlaF-(6xHis-MlaE)-MlaD(TMΔLptC TM)-MlaC-MlaB: Expression and Complementation plasmid for MlaD TM swap with LptC TM | This study |
| pBEL2138 | MlaF-(6xHis-MlaE)-MlaD(TMΔLetB TM)-MlaC-MlaB: Expression and Complementation plasmid for MlaD TM swap with LetB TM | This study |
| pBEL1198 | MlaF-MlaE-(6xHis-MlaD)-MlaC-MlaB: Expression and complementation plasmid for MlaFEDCB, with N-terminal his tag on MlaE | This study |
| pBEL2093 | MlaF-MlaE(Δ1-15)-(6xHis-MlaD)-MlaC-MlaB: Expression and Complementation plasmid for MlaE IF1 1-15 aa deletion | This study |
| pBEL2132 | MlaF-MlaE(Δ1-25)-(6xHis-MlaD)-MlaC-MlaB: Expression and Complementation plasmid for MlaE IF1 1-25 aa deletion | This study |
| pBEL2092 | MlaF-MlaE(Δ1-30)-(6xHis-MlaD)-MlaC-MlaB: Expression and Complementation plasmid for MlaE IF1 1-30 aa deletion | This study |
| pBEL2133 | MlaF-MlaE(Δ1-39)-(6xHis-MlaD)-MlaC-MlaB: Expression and Complementation plasmid for MlaE IF1 1-39 aa deletion | This study |
| pBEL2099 | MlaF-MlaE(Y81A)-(6xHis-MlaD)-MlaC-MlaB: Complementation plasmid for MlaE Y81A | This study |
| pBEL2100 | MlaF-MlaE(Y81W)-(6xHis-MlaD)-MlaC-MlaB: Complementation plasmid for MlaE Y81W | This study |
| pBEL2098 | MlaF-MlaE(R97A)-(6xHis-MlaD)-MlaC-MlaB: Complementation plasmid for MlaE R97A | This study |
| pBEL2094 | MlaF-MlaE(E98A)-(6xHis-MlaD)-MlaC-MlaB: Complementation plasmid for MlaE E98A | This study |
| pBEL2095 | MlaF-MlaE(K205A)-(6xHis-MlaD)-MlaC-MlaB: Complementation plasmid for MlaE K205A | This study |
| pBEL2097 | MlaF-MlaE(D250A)-(6xHis-MlaD)-MlaC-MlaB: Complementation plasmid for MlaE D250A | This study |
| pBEL2057 | MlaF-(6xHis-MlaE, Tyr81Bpa)-MlaD-MlaC-MlaB: Expression plasmid for MlaE Tyr81Bpa mutant for *in vivo* crosslinking | This study |
| pBEL2060 | MlaF-(6xHis-MlaE, Val77Bpa)-MlaD-MlaC-MlaB: Expression plasmid for MlaE Val77Bpa mutant for *in vivo* crosslinking | This study |
| pBEL2061 | MlaF-(6xHis-MlaE, Leu78Bpa)-MlaD-MlaC-MlaB: Expression plasmid for MlaE Leu78Bpa mutant for *in vivo* crosslinking | This study |
| pBEL2062 | MlaF-(6xHis-MlaE, Leu70Bpa)-MlaD-MlaC-MlaB: Expression plasmid for MlaE Leu70Bpa mutant for *in vivo* crosslinking | This study |
| pBEL2063 | MlaF-(6xHis-MlaE, Leu99Bpa)-MlaD-MlaC-MlaB: Expression plasmid for MlaE Leu99Bpa mutant for *in vivo* crosslinking | This study |
| pBEL2065 | MlaF-(6xHis-MlaE, Trp149Bpa)-MlaD-MlaC-MlaB: Expression plasmid for MlaE Trp149Bpa mutant for *in vivo* crosslinking | This study |
| pBEL2066 | MlaF-(6xHis-MlaE, Phe209Bpa)-MlaD-MlaC-MlaB: Expression plasmid for MlaE Phe209Bpa mutant for *in vivo* crosslinking | This study |
