## Supplemental file 2 for "Structure of bacterial phospholipid transporter MlaFEDB with substrate bound"

**Supplementary file 2: Data refinement statistics for cryo EM structure of MlaFEDB**

| Number of particles: | 177,513 |
| --- | --- |
| Final resolution (FSC = 0.143): | 3.05 Å |
| Symmetry imposed: | none |
| B-factor for sharpening: | -50 Å^2^ |
| Sphericity of 3DFSC: | 0.961 |
| Map CC (mask): | 0.8555 |
| Map CC (volume): | 0.8363 |
| Map CC (peaks): | 0.6908 |
| rmsd (bonds): | 0.003 Å |
| rmsd (angles): | 0.549 ° |
| All-atom clashscore: | 8.32 |
| Ramachandran plot values: |  |
| outliers: | 0.00 % |
| allowed: | 3.01 % |
| favored: | 96.99 % |
| Rotamer outliers: | 0.06 % |
| C-beta deviations: | 0.00 % |
| Overall score (Molprobity): | 1.62 |
| Deposition: |  |
| EMDB ID: | EMD-22116 |
| PDB ID: | 6XBD |
