## Supplemental file 1 for "Structure of bacterial phospholipid transporter MlaFEDB with substrate bound"

**Supplementary file 1: Data collection parameters for cryoEM structure of MlaFEDB**

| microscope: | Titan Krios |
| --- | --- |
| electron energy (kV): | 300 |
| defocus range (Å): | -1500 to -3500 |
| pixel size in super-resolution mode (Å): | 0.416 |
| total electron dose (e-/Å^2^): | 71 |
| number of frames in each movie: | 30 |
| frame rate (ms): | 200 |
| number of images acquired: | 3212 |
| number of particles picked: | 1,283,606 |
| number of particles selected after 2D classification: | 731,205 |
| pixel dimension of individual windows: | 300x300 (initial rounds)  500x500 (final model) |
