## Supplemental figures for "Structure of bacterial phospholipid transporter MlaFEDB with substrate bound"

#### Figure 1 — figure supplement 1

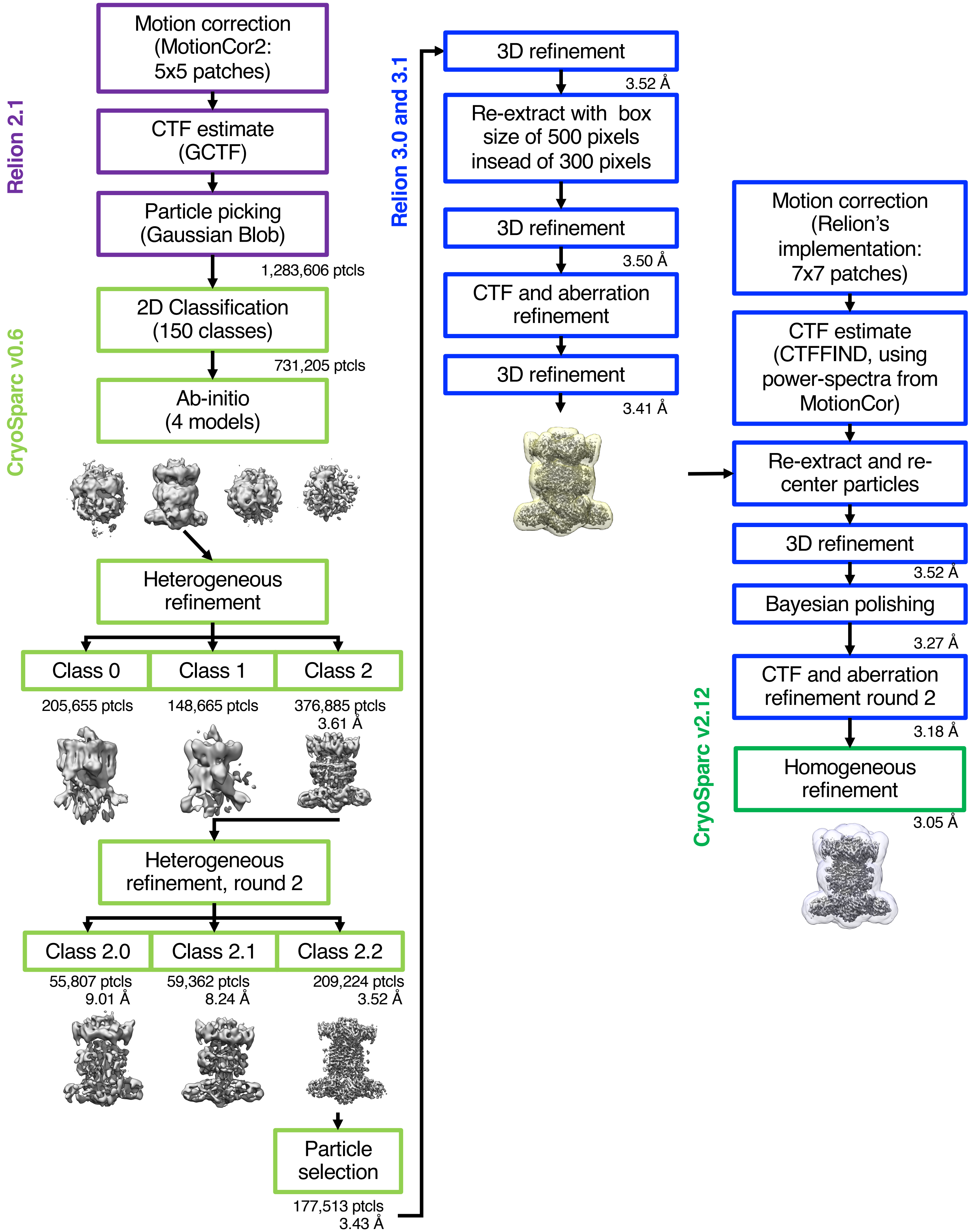


**Figure 1 — figure supplement 1. Cryo-EM data processing workflow.** In column 2, an intermediate map is shown with the mask used during reconstructions in relion, and in column 3, the map is shown with the "mask_refine" automatically computed by Cryosparc.

### Figure 1 — figure supplement 2

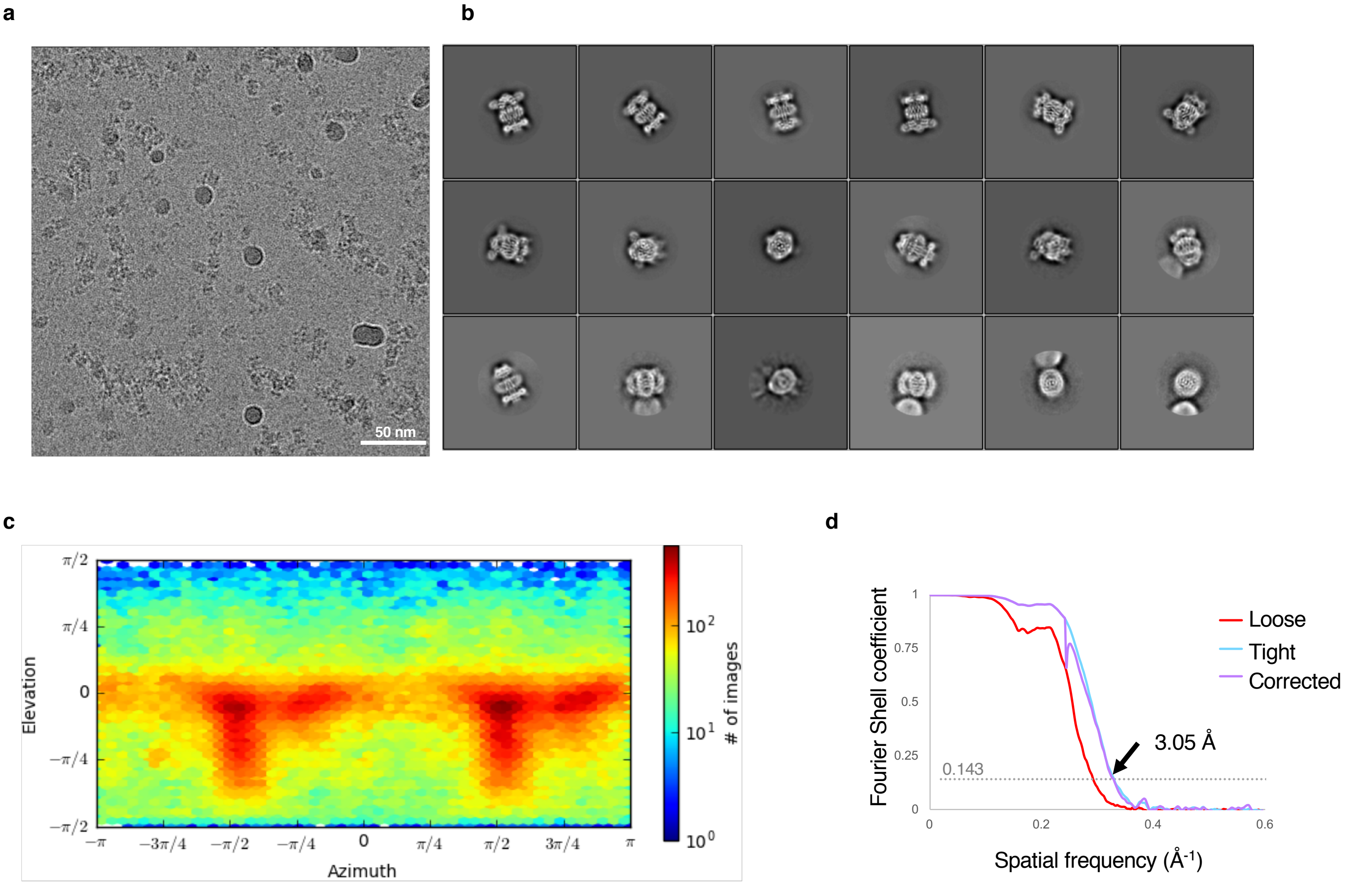


**Figure 1 — figure supplement 2. Cryo EM data and analysis. (a)** Representative micrograph. **(b)** Representative 2D classes. **(c)** Viewing direction distribution of the final model from Cryosparc. **(d)** Gold-standard FSC curve.

#### Figure 1 — figure supplement 3

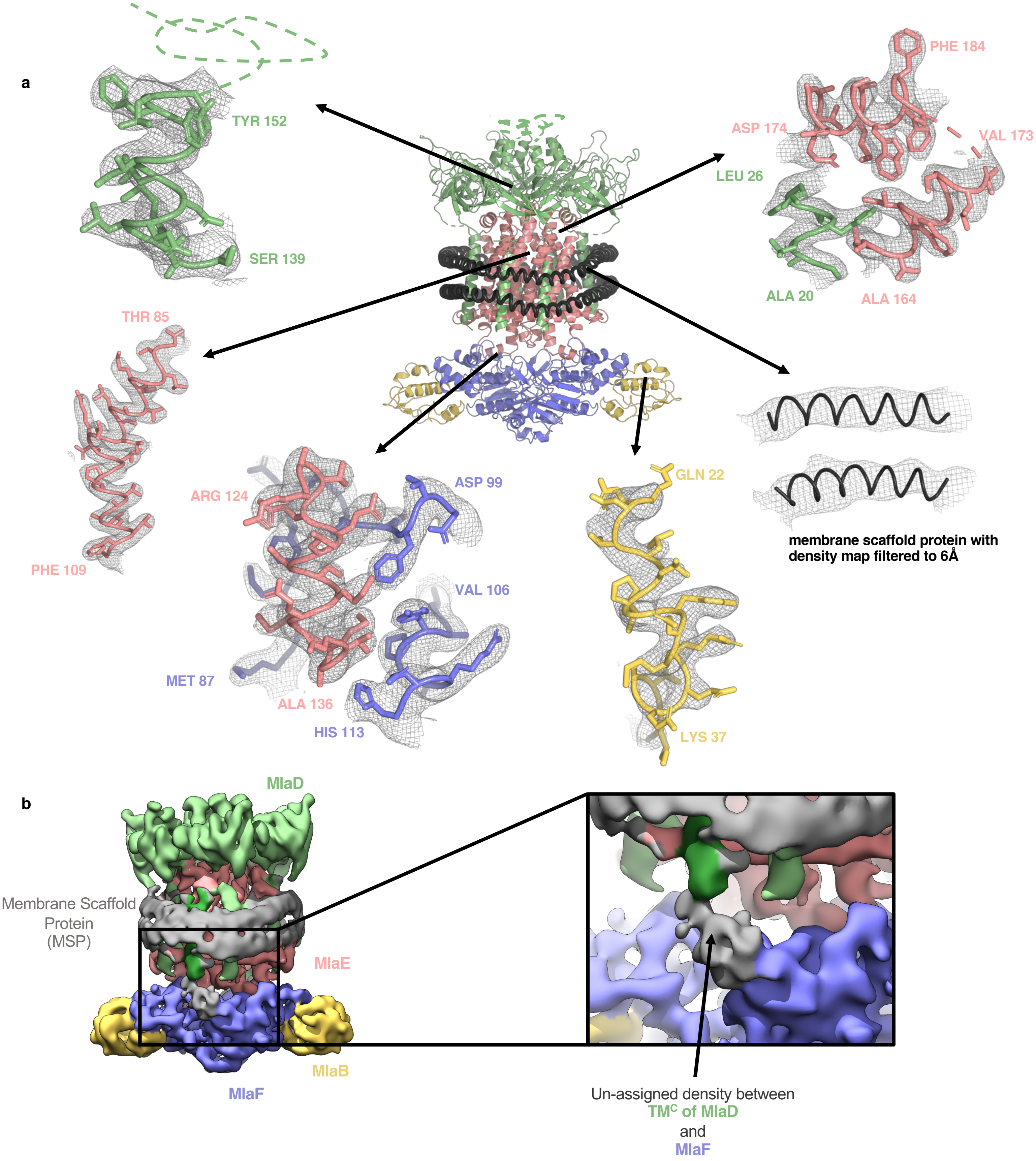


**Figure 1 — figure supplement 3. Representative density for MlaFEDB complex. (a)** Representative densities from the final map into which the model was built. **(b)** Density map filtered to 6 Å showing unmodelled density between $TM^C$ of MlaD (and $TM^F$ on the other side) and MlaF. This density is not well defined, and may correspond to the His-Tag from MlaD, or an unknown ligand.

#### Figure 1 — figure supplement 4

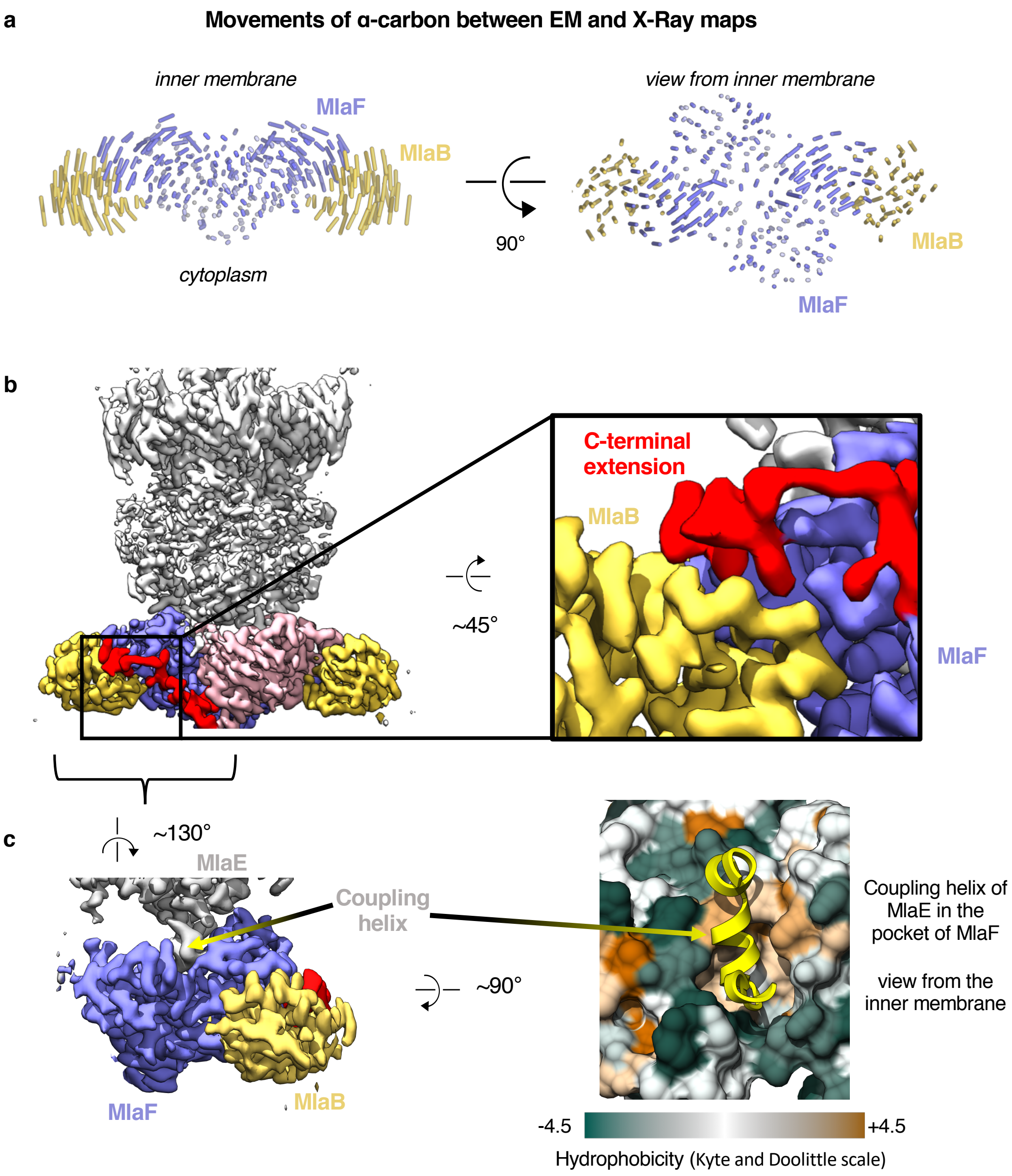


**Figure 1 — figure supplement 4. Comparison of MlaFB from X-ray and Cryo-EM structures. (a)** The EM (this study, PDB 6XBD) and X-Ray (Kolich *et al.*, 2020, PDB: 6XGY) models were aligned on the MlaF dimer, and to illustrate differences, lines between alpha-carbons are shown using a modified version of the ColorByRMSD pymol script. **(b)** Cryo-EM density map of MlaFEDB. The C-terminal extension (red) of one MlaF monomer (pink) wraps around the other MlaF monomer (slate blue), placing a small helix at the MlaF/MlaB interface, consistent with the X-Ray structure of MlaFB alone (Kolich *et al.*, 2020). **(c)** Interaction between MlaF and the coupling helix of MlaE in the cryo EM structure.

#### Figure 1 — figure supplement 5

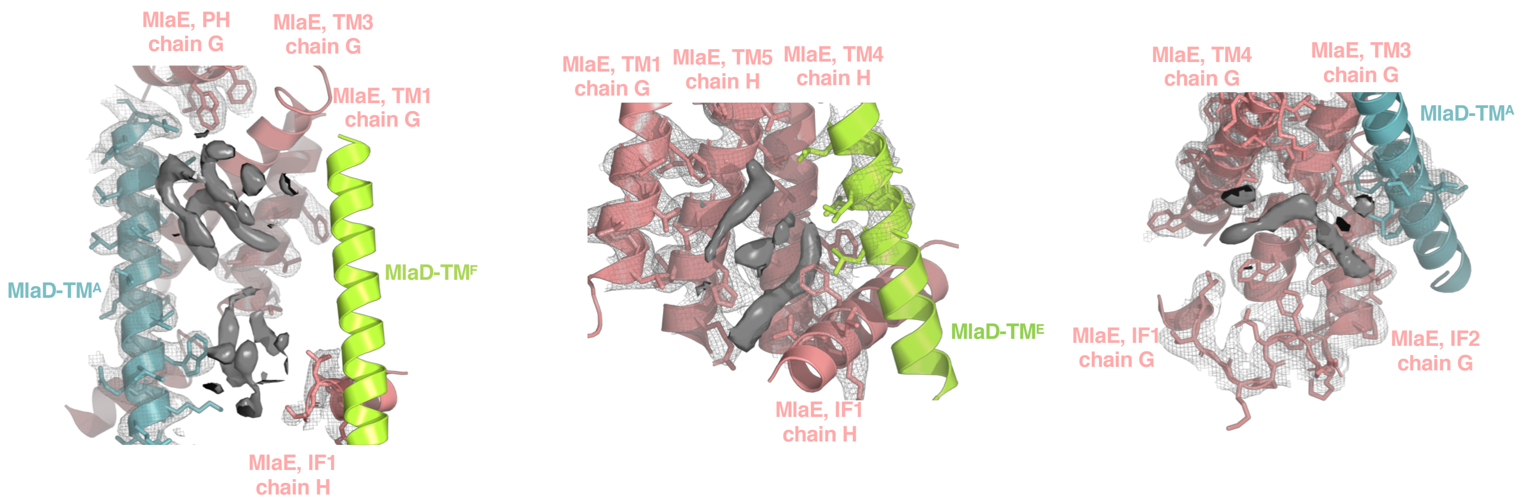


**Figure 1 — figure supplement 5. Extra densities in the transmembrane region.** Elongated densities around the TM region and in the cleft formed by IF1 are shown in solid grey, which could potentially be either phospholipids or other molecules. Due to the ambiguous nature of the densities, we have not modeled any coordinates in these regions.

#### Figure 2 — figure supplement 1

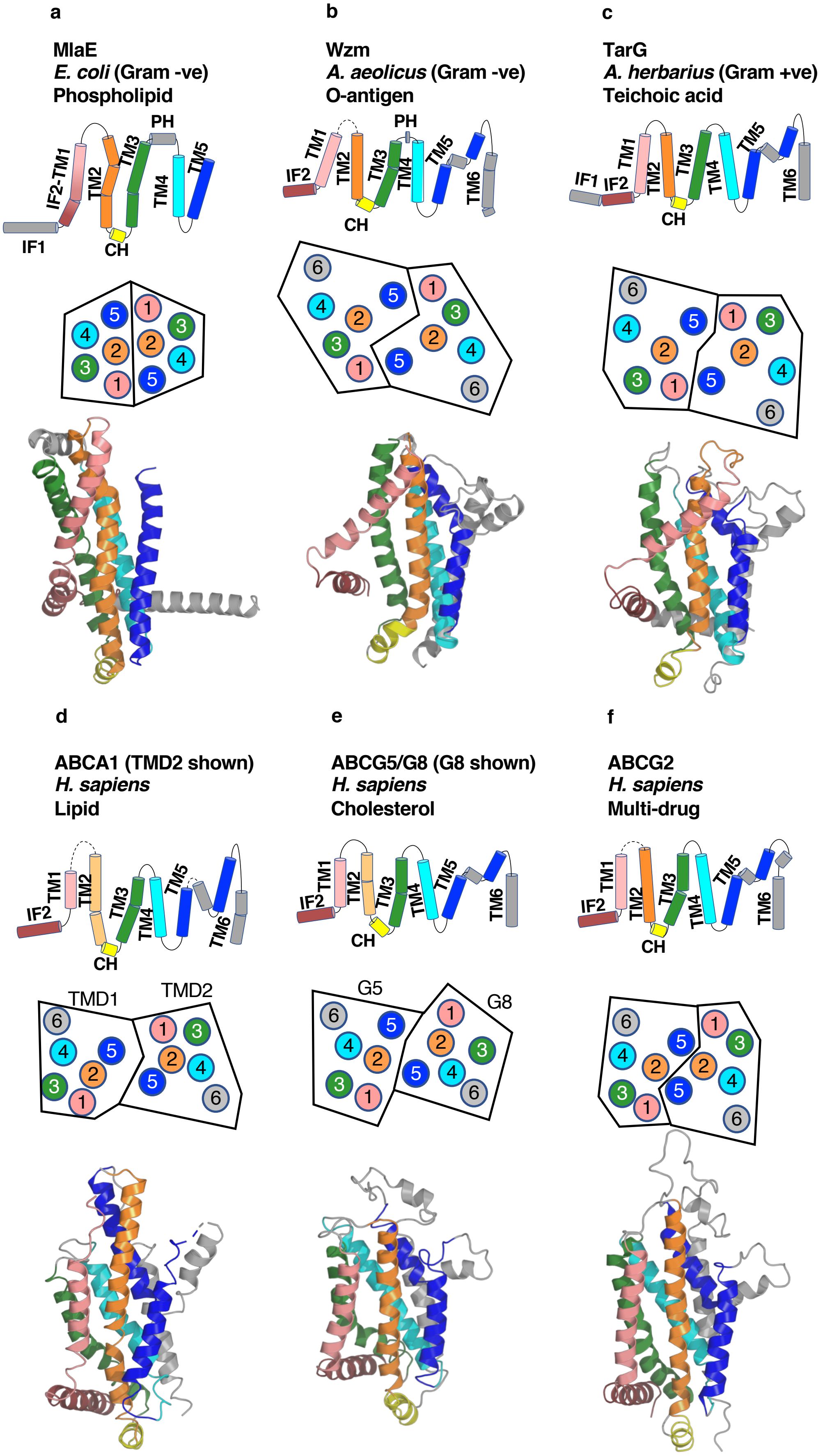


**Figure 2 — figure supplement 1. Comparison of MlaE to other ABC transporters. (a)** MlaE, this paper, PDB 6XBD; **(b)** O-antigen transporter, PDB 6OIH; **(c)** TarG, PDB 6JBH; **(d)** ABCA1, PDB 5XJY, TMD2 shown; **(e)** ABCG5/G8, PDB 5DO7, G8 shown; **(f)** ABCG2, PDB 6HBU. Topology diagrams (first row); schematics representing helices at the dimer interface, viewed from the periplasm (each circle represents a helix) (second row); and cartoon views of monomers (third row) are shown.

### Figure 2 — figure supplement 2

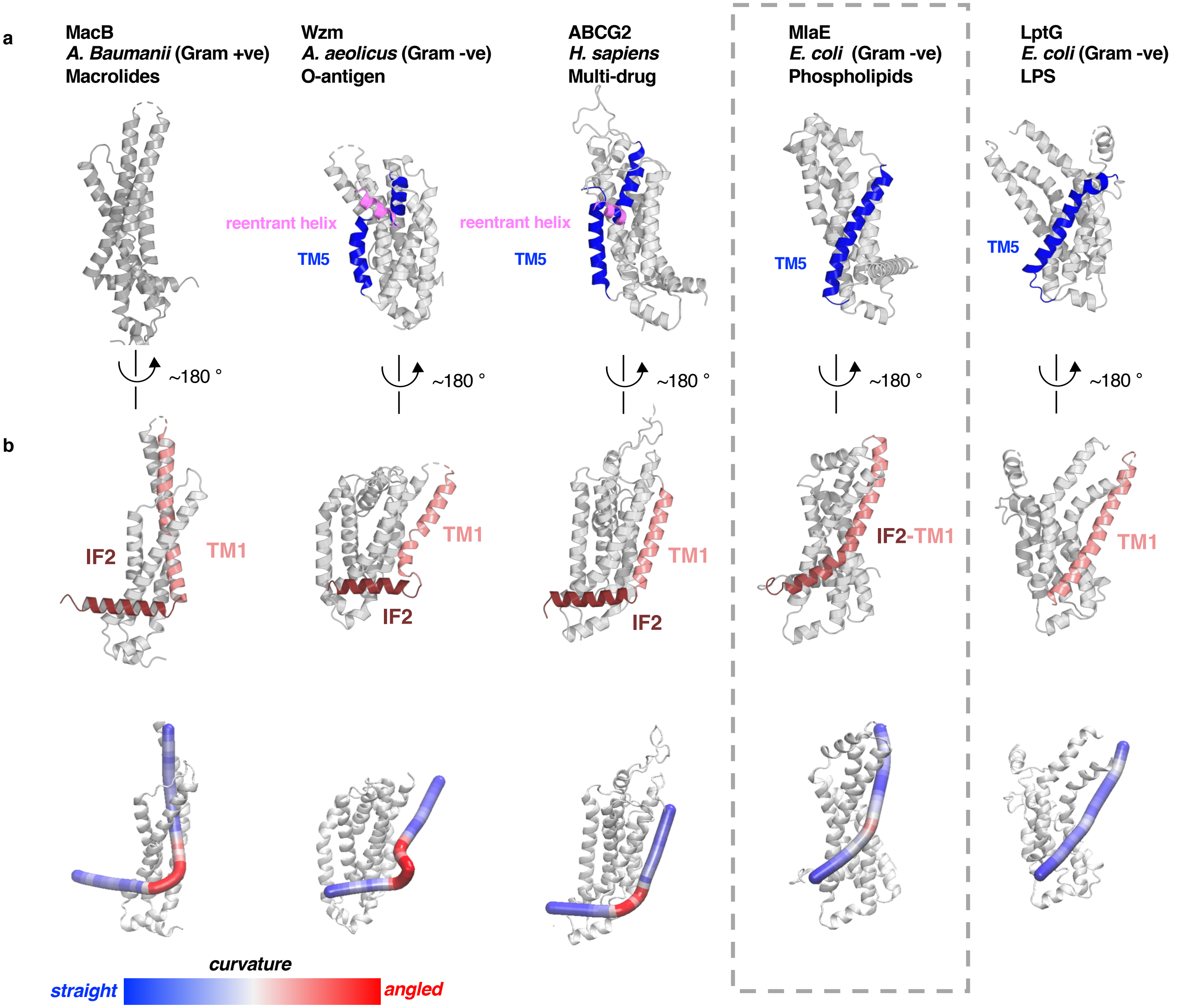


**Figure 2 — figure supplement 2. Structural variation in MlaE TMD compared to other ABC transporters. (a)** TM5 helices (blue) are either a single continuous helix (MlaE, PDB 6XBD; LptG, PDB 6MHZ), a reentrant helix insertion (pink) (Wzm, PDB 6OIH; ABCG2, PDB 6HBU), or no helix (MacB, PDB 5GKO). **(b)** IF2 (brown) and TM1 (salmon) helices adopt a range of conformations, from IF2 being parallel to the membrane interface and forming a sharp angle with TM1 (left) to one continuous smooth helix for LptG (right). Top row, cartoon representation; bottom row, curvature color coded using Bendix (Dahl, Chavent and Sansom, 2012).

#### Figure 3 — figure supplement 1

**a**

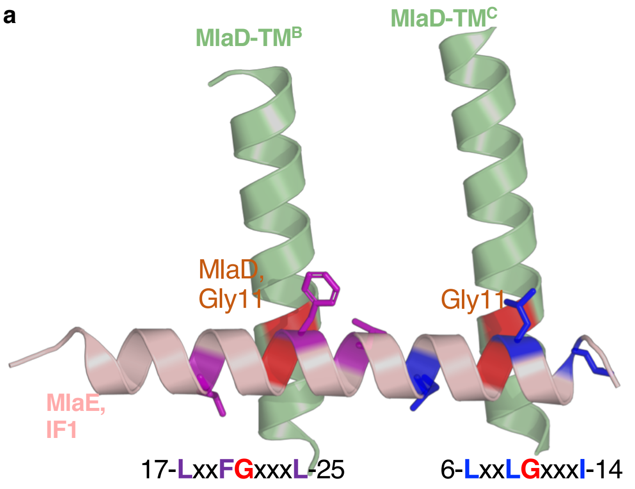


**b**

**MlaD**

```
                                                  10        20        30
                                                  |         |         |
E. coli WP_001296448.1/1-183               -MQTKKNEIWVGIFLLAALLAALFVCLKAANVTS
R. rickettsii AFB21753.1/1-149             -MQQNIIETIIGFVVL--IIALLFLIFAYKTGSS
N. moscoviensis WP_053378545.1/1-147       -MEKTKLELAVGVFVLVGVVCLGYLSIKLGKLEV
F. tularensis WP_003014807.1/1-177         -MRNKYFETSVGIFIIIGVLCLLFLTFKVSGTSF
N. gonorrhoeae WP_071197970.1/1-165        -MKKNILEFWVGLFVLIGAAAVAFLAFRVAGGAA
A. baumannii WP_000842362.1/1-226          -MKSRTSELAVGIFVIIFGIALFFLAMKVSGLVG
B. cepacia AIO25115.1/1-181                -MKKTALDFWVGLFVVVGFLAVLFLALKVGNMSS
L. pneumophila WP_027221206.1/1-158        MKKQRYVDISVGIFMLLGLLALLVMAMKVSNITD
P. luteoviolacea AOT09785.1/1-181          -MNTRKIEILVGLFVALGVAAFVMLAMKVANAGI
V. cholerae AAF95660.1/1-163               MQQTRKIEFWVGSFVIAGICAILVMIFQVADVKG
K. pneumoniae WP_002918387.1/1-183         -MQTKKTEIWVGVFLLVALLAALFVCLKAANVTS
H. influenzae P45029.1/1-167               MRQTIKYEFWVGLFLLLGIGALVFLGLRVANVQG
P. aeruginosa WP_003094339.1/1-157         -MQTRTLEIGVGLFLLAGLLALLLLALRVSGLSV
```

**c**

**MlaE**

```
                                          10        20        30        40        50        60        70        80
                                          |         |         |         |         |         |         |         |
E. coli EIQ68638.1/1-260 /1-260            ----MLLNALASLGHKGIKTLRTFGRAGLMLFNALVGKPEFRKHAPLLVRQLYNVGVLSMLIIVVSGVFIGMVLGLQGYLVLTTYSAETSLGM
R. rickettsii AFB22698.1/1-259            ----MLFNIANSVGKRTVKFAQSVGSFSLFSFAAVSSIIRPPLYLSLIIRQLLFIGFHSLPVVAMTTFFSGAVLALQSYIGFSRFSAESSIAT
N. moscoviensis ALA57157.1/1-256          -----MIHLIERIGAFVLKYVGEMGRMLIFVLSAFAWLIRPPLRFAQIMKQVHFIGYKSTFVVVLTAGFTGMVLGLQGYYTLRKFGSEALLGS
F. tularensis WP 003035471.1/1-254        ------MTIFSKVYTLTFSLIYSLVKALSLILRIVISKLNIRD----CIIQIKAVGVNSIIIIVTSGVFIGLVLSLQGYYTLSKFGAHSLLGV
N. gonorrhoeae WP 071197969.1/1-258       ------MNFIRSVGAKTLGFIQSLGSITLFLLNILAKSGTAFVRPRLSVRQVYFAGVLSVLIVAVSGLFVGMVLGLQGYTQLSKFKSADILGY
A. baumannii WP 001091949.1/1-258         ------MNTIAWLGRLVIERIRGIGVAALMLLQIIFSLPSAG-GFGRFVYQMHRVGVMSLLIITVSGLFIGLVLGLQGYSILVNVGSESMLGT
B. cepacia AIO25368.1/1-255               --------MISAIGRYVIGGLERAGYGTRLFVRLVLEFFPLLRRPRLVTKQIHFLGNYSFVIIAVSGLFVGFVLGLQGYYTLNRYGSEQALGL
L. pneumophila AMP90450.1/1-260           -----MLDTIARLGNRGVRALQGIGSSGLFLFSMLFRKPNVLRLWPLVRYQLYFIGVLSCLIIVVSALFIGMVVGLQGYNTLQKFGASSQLGQ
P. luteoviolacea AOT09786.1/1-258        ------MDFLQKLGRKTLNRFASIGRATQMLLGALLNVPNLRQGTPLLIKQLYMVGHQSLLIIMVSGLFIGMVLALQGYTVLVGYGAEDSLGP
V. cholerae AAF95661.1/1-258              ------MRWVSSVGQRTLAVSETFGRASLMLFGALVGRPQPIRHFPLLVRQLYSIGVQSLAIIIVSGLFIGMVLSLQGYVILVDYGAETSLGQ
K. pneumoniae WP 004150949.1/1-260        ----MLFNALAALGHRGIKTTATFGRAGLMLFNAVVGKPEFRKHAPLLVRQLYNVGVLSMLIIIVSGLFIGMVLGLQGYLVLTTYSAETSLGM
H. influenzae AAC22742.1/1-261            ----MIVNFISALGKQVIDFFRALGRAGFMLFGALIGKPQIRKHFPLLVKQMHVLGVQSLLIILLSGLFIGMVLGLQGYVVLIDFSAETSLGQ
P. aeruginosa KEA30503.1/1-265            MRRVSPLERIRLFGRAGLDVVAALGRSTLFLGHALLGRRTPGTGLHLLVKQLYSVGVLSLAIIVVSGLFIGMVLALQGYNILISYGSEQAVGQ

                                          90        100       110       120       130       140       150       160       170       180
                                          |         |         |         |         |         |         |         |         |         |
E. coli EIQ68638.1/1-260 /1-260            LVALSLLRELGPVVAALLFAGRAGSALTAEIGLMRATEQLSSMEMMAVDPLRRVISPRFWAGVISLPLLTVIFVAVGIWGGSLVGVSWKGIDS
R. rickettsii AFB22698.1/1-259            VVVLSLTRELGPVLAGLMVAGRVGASIAAEIATMRVTEQVDALYTLSTDPIKYLVFPRVITAIITMPCLVLIGDIIGVMGGYLVGVYKLDFNS
N. moscoviensis ALA57157.1/1-256          AVALSMIRELGPVLAALMVTARAGSAMTAEIGIMRITEQIDALDTMAVNPLQYLIAPKLVAGLIGVPLLVAIFDVVGIYGGYLVGVDLLGVSA
F. tularensis WP 003035471.1/1-254        MVALSVLRELGPVVTAMLFAGRACSSITSEIGLMKATDQINSLKVMNVNPISFILSTRFWACMISGPILALIFASVAILAGFILAEAALGISY
N. gonorrhoeae WP 071197969.1/1-258       MVAASLLRELGPVLAAILFASSAGGAMTSEIGLMKTTEQLEAMNVMAVNPVARVVAPRFWAGVFSMPLLASIFNVAGIFGAYLVGVSWLGLDS
A. baumannii WP 001091949.1/1-258         MVSLTLLRELAPVVAALLFAGRAGSALTAEIGSMKQSEQLASMEMIGVDPLKQIVSPRLWAGIVSLPMLTVIFAAIGIVGGKLVGVDFLGVDE
B. cepacia AIO25368.1/1-255               LVALSLVRELGPVVTALLFAGRAGTSLTAEIGLMKAGEQLTALEMMAVDPIKNVIAPRLWAGIIAMPLLAAIFNAVGVLGGYFVGVVLIGVDP
L. pneumophila AMP90450.1/1-260           LLALSVARELGPVISALLFAGRAGSALTAEIGLMKATEQLSSMDMMGVDPLGRVVYPRFVAGFIALPILALIFSAVAIYGGYFIGVYWLGVDA
P. luteoviolacea AOT09786.1/1-258        LVALSLLRELGPVVTALLFAGRAGSALTAEIGLMKATEQLSSLEMMAVDPLKRVVAPRFWAGFISMPLLALIFSAVAILGAHLVGVDWLGVDS
V. cholerae AAF95661.1/1-258              MVALSLLRELGPVVTALLFAGRAGSALTAEIGLMKATEQLSSLEMMAVDPLKRVIAPRFWAGVISMPLLAMIFMAVGIWGGQLVGVDWKGIDH
K. pneumoniae WP 004150949.1/1-260        LVALSLLRELGPVVAALLFAGRAGSALTAEIGLMRATEQLSSMEMMAVDPLRRVISPRFWAGVISLPLLTIIFVAVGIWGGALVGVSWKGIDG
H. influenzae AAC22742.1/1-261            LVALSLLRELGPVVTALLFAGRAGSALTAEIGLMKATEQLSSLEMMAVDPLRRVIAPRFWAGVISMPVLSILFIAIGIWGGSLVGVDWKGVDS
P. aeruginosa KEA30503.1/1-265            MVALTLLRELGPVVTGLLFAGPAGSALTAEIGNMKATEQLSSLEMIGVDPLKYIVAPRLWAGFISMPLLAAIFSVVGIWGGAMVAVDWLGVYE

                                                     190       200       210       220       230       240       250       260
                                                     |         |         |         |         |         |         |         |
E. coli EIQ68638.1/1-260 /1-260            GFFWSAMQNAVDWRMDLVNCLIKSVVFAITVTWISLFNGYDAIPTSAGISRATTRTVVHSSLAVLGLDFVLTALMFGN-
R. rickettsii AFB22698.1/1-259            AAYLTSTFQYLE-PIDVISGLVKAGVFGFIISIISCYSGYYSGKGAKGVGRATTSAVVNSSILILISNYLITELFFKV-
N. moscoviensis ALA57157.1/1-256          GSYWTSIESAVEWK-DVYGGILKSVSFGLIVSWVCCYKGFYTRMSAEGLGTATTEAVVLSSVLILVWDYFLTSVLL---
F. tularensis WP 003035471.1/1-254        GEFWSNIQSSVT-ASDISNGIIKSIVFAFITAWIALYQGYYCIPDSNGIAKATTKTVVYCCMSVLGADLILTSIMFGGV
N. gonorrhoeae WP 071197969.1/1-258       GIFWSQMQNNITIHYDVINGLIKSAAFGVAVTLIAVHQGFHCVPTSEGILRASTRTVVSSALTILAVDFILTAWMFTD-
A. baumannii WP 001091949.1/1-258         GSFWSGMQNNVQFGHDVVNGIIKSIVFALLCTWIAVFQGYACDPTPEGIATAMTRTVVYSSLCVLGFDFVLTAVMFGGI
B. cepacia AIO25368.1/1-255               GAFWSQMQGGVQVWADVGNGVIKSIVFGFAVTFIALFQGYEAKPTPEGVSRATTKTVVFASLAVLGLDFLLTALMFS--
L. pneumophila AMP90450.1/1-260           GSFWSNMQAAVNFRIDILSGIIKSLVFAFVVTWISVFQGFECVPTAEGISHATTKTVVYSSLAVLGLDFLLTAMMIGDW
P. luteoviolacea AOT09786.1/1-258        GSFWSIMQSQVSLQQDILNGLIKSFVFALIVTWIALYKGYDCVPTSEGISKATTETVVHSSLAVLGFDFVLTAVMFSS-
V. cholerae AAF95661.1/1-258              GSFWSAMQASVELGQDIGNSTIKCVVFAFTVTWIALFNGYDAIPTSEGISRATTRTVVHSSLAVLGLDFVLTALMFGN-
K. pneumoniae WP 004150949.1/1-260        GFFWTAMQNAVDWRMDLVNCLIKSLVFAITVTWIALFNGYDAIPTSAGISRATTRTVVHASLAVLGLDFVLTALMFGN-
H. influenzae AAC22742.1/1-261            GSFWSVMQNSVSWSYDILNGFIKAVFFAVAVTWIALFNGYDCMPTSEGISQATTRTVVHASLVVLGLDFILTAIMFGAG
P. aeruginosa KEA30503.1/1-265            GSFWANMQNSVQFTEDVLNGVIKSIVFAFVVTWIAVYQGYDCEPTSEGISRATTRTVVYASLAVLGLDFILTALMFGDF
```

**Figure 3 — figure supplement 1. Sequence conservation in MlaD and MlaE. (a)** Conserved motifs are depicted at the interface between MlaE IF1 and the TM helices of MlaD. **(b)** Sequence alignment of MlaD (TM helix only) and (**c**) MlaE from 13 diverse bacterial species**.** Red, conserved glycine residues that are part of the interaction between IF1 and the TM helices of MlaD. Blue, the conserved residues that are part of the salt bridge and polar interactions around R97. Residue numbers correspond to *E. coli* sequences. Sequence alignments were done with MUSCLE (Edgar, 2004) implemented within Phenix.

#### Figure 3 — figure supplement 2

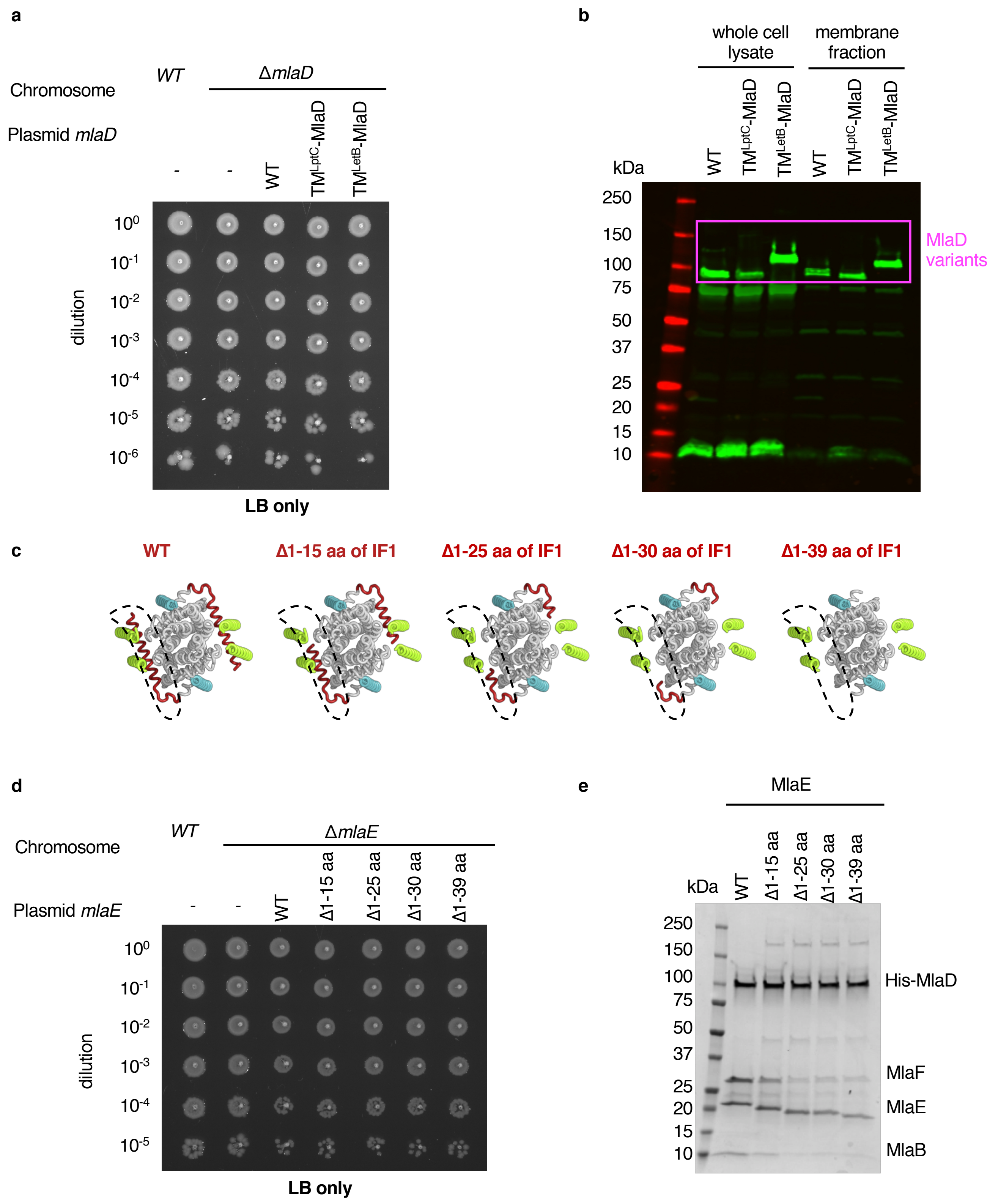


**Figure 3 — figure supplement 2. Extended data for MlaE and MlaD mutants. (a)** 10-fold serial dilutions of the indicated cultures spotted on LB agar and incubated overnight (control for data represented in **Figure 3c**). **(b)** Western blot using an anti-MlaD antibody against wild type and mutant forms of MlaD showing expression levels in the cell lysate and membrane fraction. **(c)** Illustration of the IF1 truncation mutants used in genetic complementation assays (related to **Figure 3e**). **(d)** 10-fold serial dilutions of the indicated cultures spotted on LB agar and incubated overnight (control for data represented in **Figure 3d**). **(e)** SDS-PAGE of recombinantly expressed and co-purified MlaFEDB complexes with IF1 mutants (**Figure 3e**), purified by pulling down His-MlaD.

#### Figure 4 — figure supplement 1

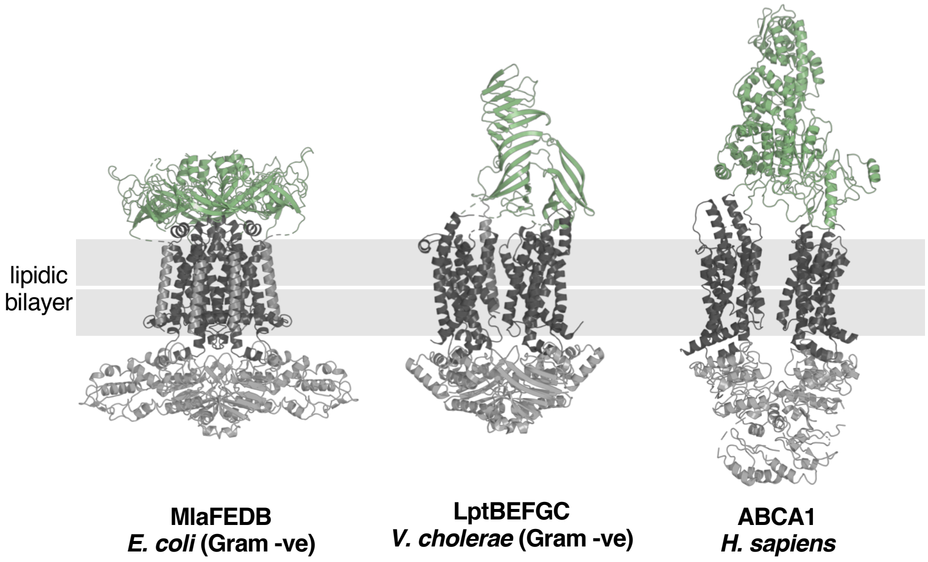


**Figure 4 — figure supplement 1. Structural diversity of extracytoplasmic lipid transport domains.** Extracytoplasmic lipid transport domains (green) from ABC transporters. MlaFEDB, PDB 6XBD; LptBEFGC, PDB 6MJP; ABCA1, PDB 5XJY.

### Figure 4 — figure supplement 2

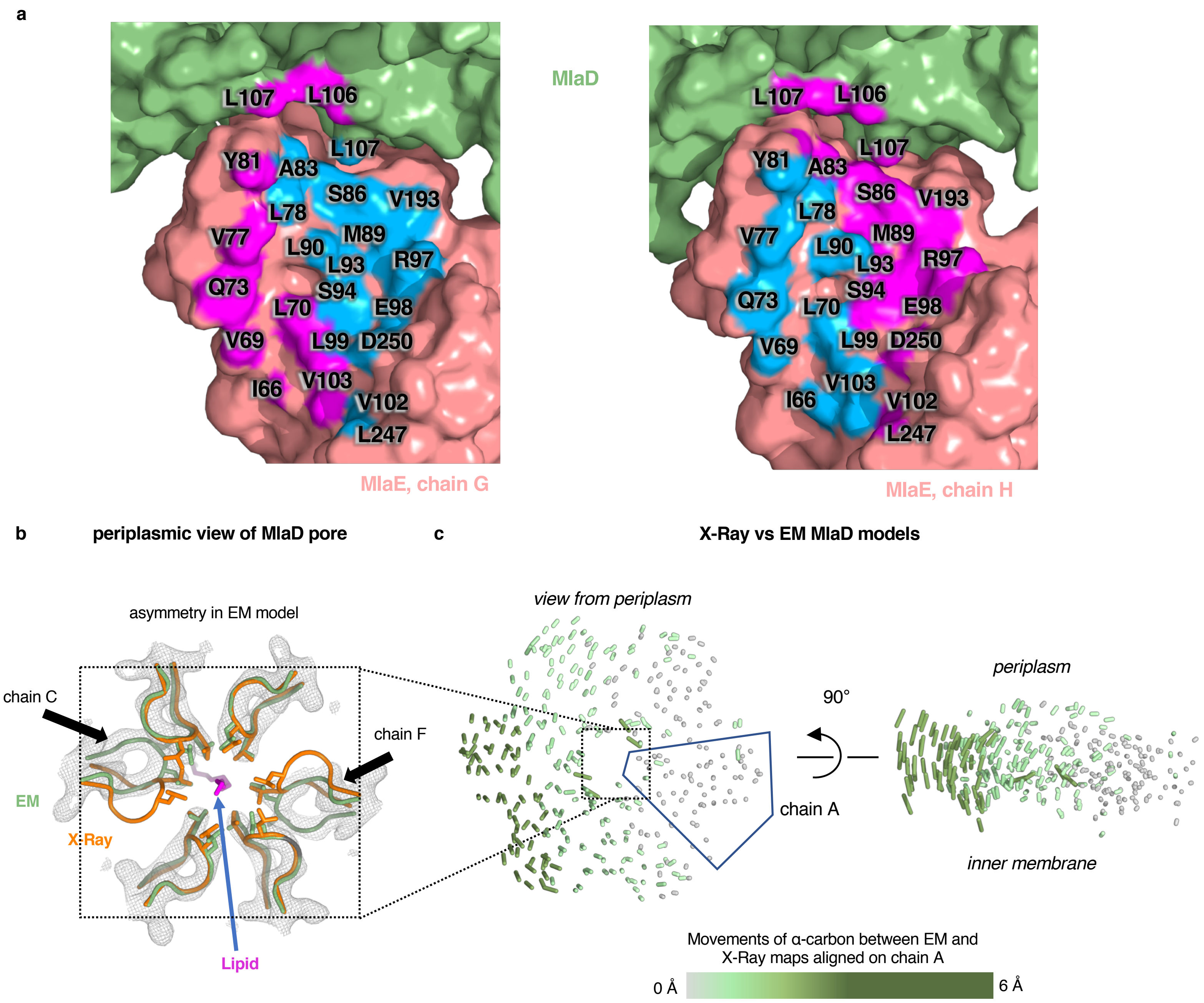


**Figure 4 — figure supplement 2. Asymmetry of lipid interactions and asymmetry of MlaD in the MlaFEDB complex. (a)** MlaE protomers (salmon) and nearby MlaD subunits (green) shown as a surface, with residues within 4.5 Å of lipid 1 colored blue and residues within 4.5 Å of lipid 2 colored magenta. **(b)** X-ray (orange, PDB 5UW2) and EM (green, PDB 6XBD) structures aligned on the whole hexamer, highlighting differences in the conformations of the pore lining loops of MlaD between the symmetric crystal structure and the asymmetric EM structure. **(c)** The EM and X-Ray MlaD structures were aligned on chain A and, to illustrate the tilt and break in symmetry, lines were drawn between alpha-carbons using a modified version of the ColorByRMSD pymol script.

#### Figure 4 — figure supplement 3

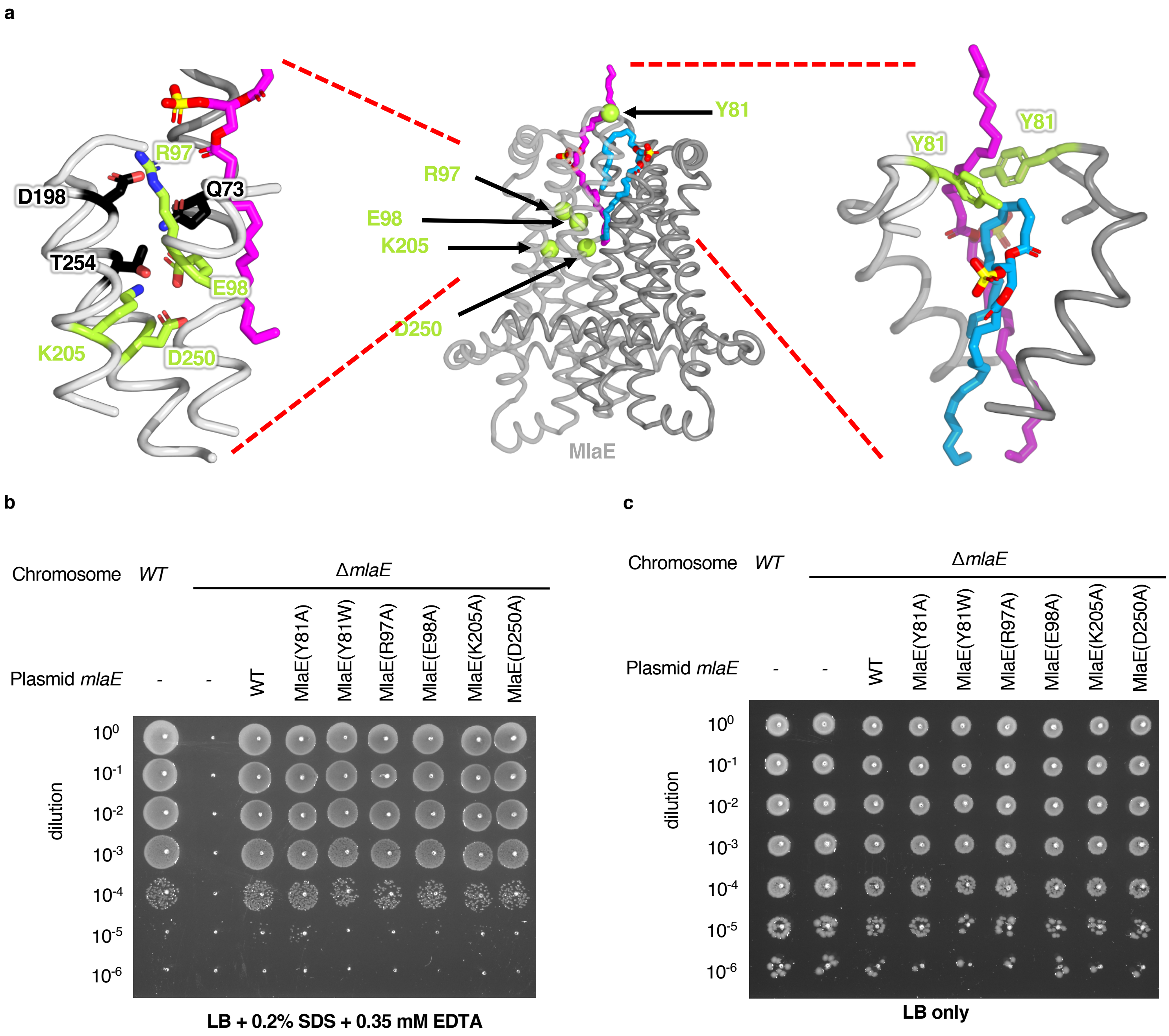


**Figure 4 — figure supplement 3. Point mutations in MlaE. (a)** Center, MlaE dimer (gray ribbons) with sites of mutations indicated by spheres (lime), and the two bound lipids (magenta and blue sticks). Left, zoomed in view of the salt bridges formed by residues Arg97, Glu98, Lys205, and Asp250 (lime sticks), part of a larger polar interaction network including Gln73, Asp198, and Thr254 (black sticks). Right, zoomed in view of interactions between Tyr81 (lime) and bound lipids (blue and magenta). **(b)** 10-fold serial dilutions of the indicated cultures spotted on LB plates containing SDS and EDTA at the concentrations indicated and incubated overnight. The *mlaE* knockout grows poorly in the presence of SDS+EDTA, but can be rescued by the expression of WT MlaE or MlaE mutants. **(c)** Same as in (b), but plated on LB only.

#### Figure 4 — figure supplement 4

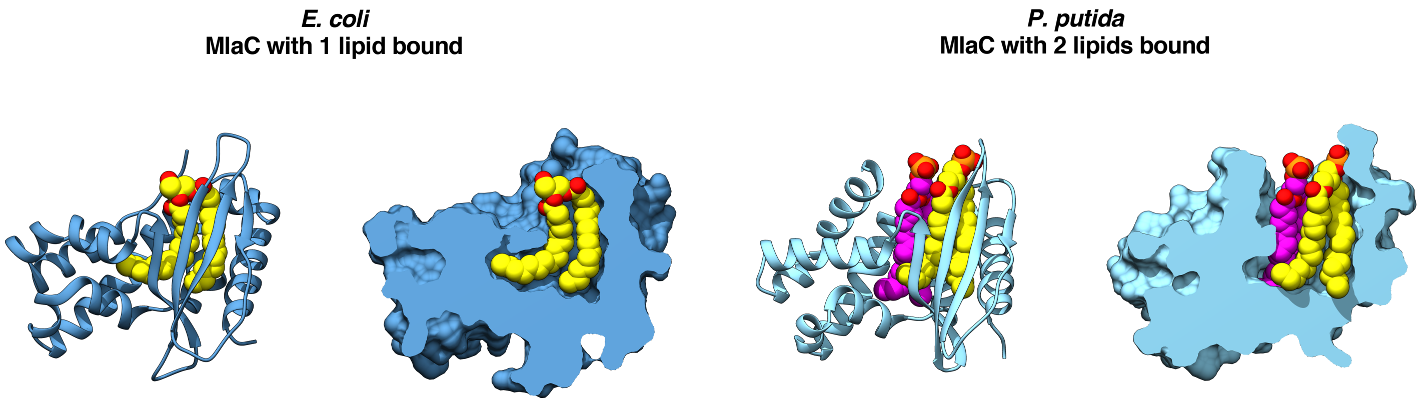


**Figure 4 — figure supplement 4. MlaC structures in which either one or two lipids are bound.** *E. coli* MlaC (PDB 5UWA) and *P. putida* MlaC (PDB 5UWB) are shown in cartoon representation and with a cross-section through the surface. Lipids are shown as yellow or magenta spheres.

#### Figure 5 — figure supplement 1

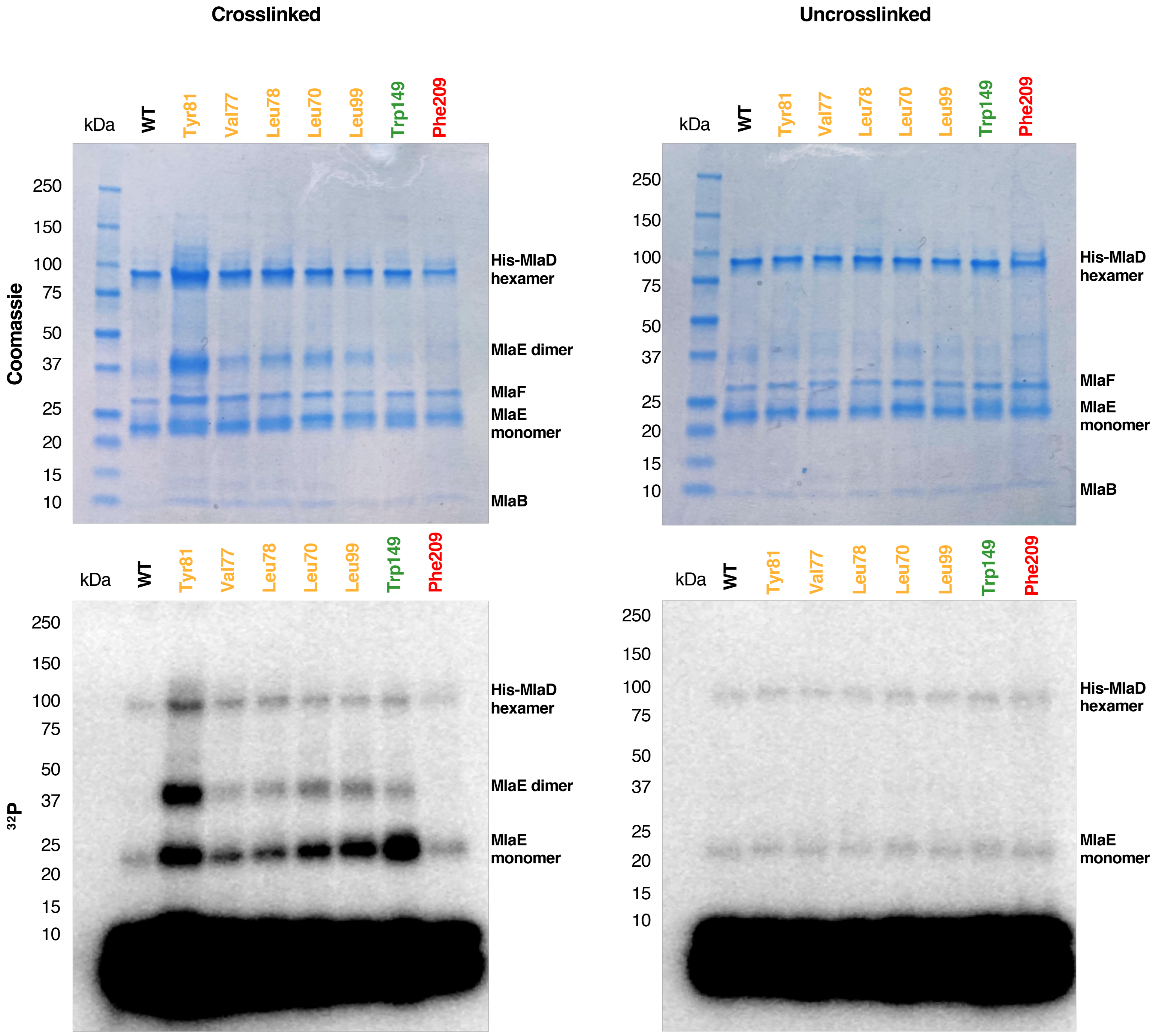


**Figure 5 — figure supplement 1. Uncropped gels of the *in vivo* photocrosslinking assay shown in Figure 5.**

### Figure 5 — figure supplement 2

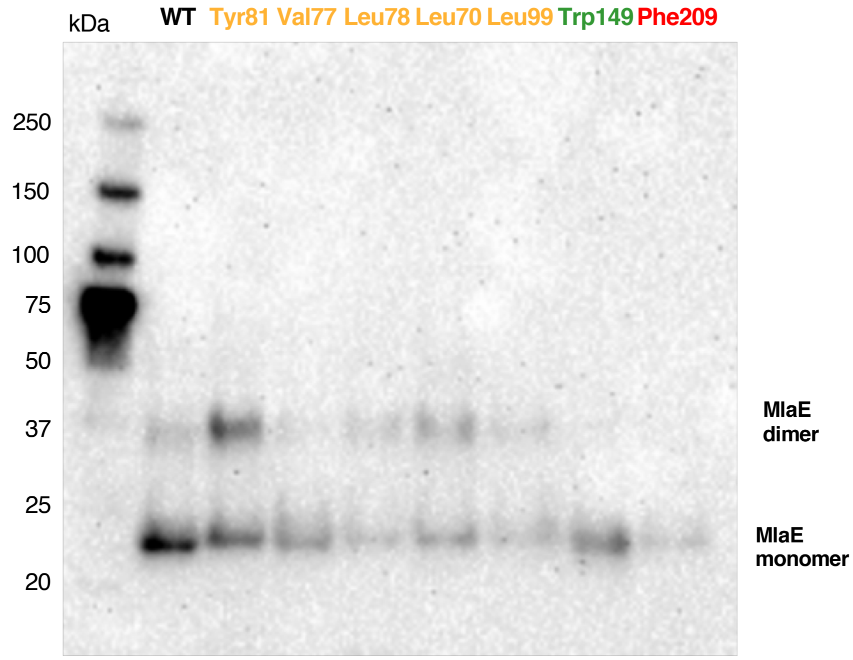


**Figure 5 — figure supplement 2. Western blot against MlaE.** Anti-His Western blot against crosslinked MlaE WT and BPA mutants.
